# Maternal high fat diet and acute viral mimic exposure impact placental inflammation, lipid peroxidation and cellular proliferation-to-death ratio across mouse pregnancy

**DOI:** 10.1101/2025.10.23.684131

**Authors:** Thaina Ferraz, Lucas Cardoso, Sadra Mohammadkhani, Enrrico Bloise, Kristin L Connor

## Abstract

Maternal obesity and viral infection induce placental inflammation, but how their co-exposure influence fetoplacental development remains unclear. We hypothesized that maternal high fat (HF) diet and viral infection would independently induce placental inflammation and lipid peroxidation, reduce antioxidant defence, and cellular turnover. Further, HF diet would compromise placental capacity to adapt to infection. Female C57BL/6J mice were fed a control (CON) or 62% HF diet six weeks before and throughout pregnancy and injected with poly(I:C) (viral mimic) or vehicle (VEH) 24h before sacrifice at gestational days (GD) 12.5, 15.5, and 18.5 (n=5–8/group/GD). Placental inflammasome (NLRP3), oxidative stress (4-HNE), antioxidant defence (GPx-4), and cellular proliferation-to-death ratio (Ki-67, Caspase-3) were assessed by immunohistochemistry, and mRNA expression of *Tlr3, Irf3, Tlr4, Tirap*, and *Il-1β* were measured by qPCR. Data were analysed by linear mixed models (p≤0.05). At GD12.5, infection was associated with increased Tlr3 mRNA and immunoreactive (ir)-4-HNE, and reduced ir-GPx-4 expression in the placental labyrinth zone (LZ). By GD15.5, HF diet was associated with increased ir-NLRP3 in both LZ and junctional zones (JZ). Exposure to infection alone and co-exposure to HF diet and infection further increased LZ ir-NLRP3. At GD18.5, HF diet was associated with increased *Tirap* and *Il-1β* mRNA expression, ir-4-HNE in the JZ and ir-Caspase-3 in the LZ. Maternal HF diet and infection exert distinct effects on the placenta across gestation, suggesting that maternal overnutrition might reduce the placenta’s capacity to handle adverse exposures, which may increase susceptibility to poor fetal outcomes.

## Introduction

Among adults of reproductive age globally, the prevalence of obesity is higher in women than men, increasing risk for obesity during pregnancy^1^. Maternal obesity is associated with altered placental development and fetal growth^2^. Poor fetoplacental growth remains a major concern, as it increases the risk of adverse outcomes at birth and predisposes offspring to long-term health complications^2^. Acute viral infections are also emerging as a critical issue during the prenatal period. For example, in Canada, between 2020 and 2021, a total of 6012 pregnant people were tested positive for SARS-CoV-2^3^. Additionally, annually, influenza affects 11% of pregnant women^4^. Although the prevalence of Zika virus in Canada is mainly restricted to travel associated infections^5^, in Brazil, more than 100,000 cases have been reported since 2016^6^ and from 2015 to 2020, 3591 cases of congenital defects were confirmed^7^.

The immunological adaptations in pregnancy required to support proper fetal development^8^ predispose women to more severe outcomes during or following viral infection compared to non-pregnant women^9^. Common viral pathogens such as influenza, Zika virus and more recently SARS-CoV-2, have been associated with increased maternal morbidity, preterm birth, and impaired fetal development^10-12^. More specifically, *in utero* exposure to Zika virus has been associated with a range of adverse offspring outcomes, from microcephaly to subtler neurodevelopmental impairments that may emerge later in childhood, even among infants without apparent birth defects at delivery^13^. Other viruses, such as cytomegalovirus, chikungunya, dengue, H1N1, herpes simplex virus, rubella and varicella zoster virus may promote significant detrimental effects in pregnancy outcome^14^.

While the mechanisms underlying these outcomes remain unclear, both obesity and viral infection during pregnancy independently induce systemic inflammation and oxidative stress, characterised by elevated circulating pro-inflammatory cytokines and reactive oxygen species (ROS), respectively^15,16^. Further, there is growing evidence that maternal obesity and viral infection are not independent challenges during pregnancy but frequently co-exist^17^. Although obesity was once most common in high-income countries, it is now increasingly prevalent in low- and middle-income countries where populations face socioeconomic disadvantage, food and nutrition insecurity, and limited access to healthcare, conditions that also increase vulnerability to infectious disease^18,3^. Such co-occurrence may contribute to poorer outcomes for both mothers and offspring^17^. Given the prevalence of obesity and viral infections in pregnancy, and that a common inflammatory mechanism may underly their independent adverse effects on the pregnancy and fetus, it is important to understand how these conditions impact fetal development so that strategies can be developed to mitigate poor fetoplacental growth and minimise the short- and long-term programming effects of these conditions on lifelong offspring health.

The placenta is a key target to understand the relationships between maternal obesity, viral infection, and pregnancy/fetal outcomes, as it serves as both a protective barrier capable of responding to infective stimuli and an exchange interface between the mother and fetus^19^. Indeed, exposure to obesity or viral infection during pregnancy is associated with increased levels of pro-inflammatory cytokines^20^ in the maternal circulation and the placenta^21^, especially in early pregnancy when physiological inflammation plays an important role in mediating implantation^22^ and placentation^23^, rendering the placenta particularly sensitive to infective insults. In fact, placenta from *in utero* deceased fetuses exhibited increase expression of pro-inflammatory markers, underlining how important placental inflammatory status is for pregnancy outcome^24^. An exacerbated *in utero* inflammatory environment can also lead to altered placental villous architecture^25,26^, impaired vascularisation^27,18^, and disrupted nutrient transporter expression and activity^27,28^. These placental structural and functional changes can impair oxygen and nutrient delivery to the fetus and may explain poor offspring outcomes observed following exposure to obesity or viral infection^21,29^.

In pregnancies where the mother has obesity, one proposed mechanism to explain altered fetoplacental development involves upregulation of placental toll-like receptor 4 (TLR4)^30^, which is activated by damage-associated molecular patterns (DAMPs) such as fatty acids^31^. TLR4 signalling through adaptor proteins like TIR domain-containing adaptor protein (TIRAP), can trigger pro-inflammatory cytokine production, including interleukin-1β (IL-1β)^32^, and a signalling cascade that will culminate in cell death^33^. Conversely, viral infections typically activate other TLRs, particularly TLR3, which recognises double stranded RNA (dsRNA), and single-stranded viruses that generate dsRNA intermediates during replication (e.g., influenza, SARS-CoV-2)^34^. Recognition of dsRNA during viral replication triggers downstream signal transduction through the IL-1 receptor domain-containing adapter-inducing interferon β (TRIF) and interferon regulatory factor 3 (IRF3) pathway^35^. This process also upregulates the expression of pro-inflammatory cytokines such as IL-1β and NLR family pyrin domain containing 3 protein, a component of the NLRP3 inflammasome^35^.

Sustained placental inflammation, induced by either obesity or infection, may disrupt the balance between ROS production and antioxidant defences, leading to oxidative stress^7^. Increased levels of oxidative stress are marked by oxidized lipids, known as lipid peroxidation^36^, where a key end product is 4-Hydroxynonenal (4-HNE)^36,37^. In high concentrations, 4-HNE can affect placental cellular dynamics, characterised by alterations in cell proliferation and cell death and further increase trophoblast production of pro-inflammatory cytokines^32^. In healthy pregnancies, placental cells can neutralise the activity of 4-HNE through the production of an antioxidant enzyme, glutathione peroxidase-4 (GPx4)^38^. Under adverse conditions in humans, a pro-inflammatory placental environment, marked by increased IL-1β, is associated with lower GPx4 expression and activity^38^ during the first trimester^39^, reducing placental capacity to neutralise 4-HNE, and thus, impedes placental protection from lipid peroxidation^38^.

Collectively, existing evidence indicates that inflammatory and oxidative stress pathways are important regulators of placental cellular dynamics, structure, and development during pregnancy^40,41^. These dynamic changes throughout gestation highlight that is important to understand how the placenta adapts to these metabolic and infectious stressors across pregnancy to maintain fetal development. When obesity and viral infection coexist, their simultaneous activation of inflammatory and oxidative pathways may amplify placental stress and worsen fetoplacental outcomes, but the ultimate phenotype likely depends on timing of exposure. However, we have poor understanding of how gestational stage informs placental sensitivity to inflammatory and oxidative stress insults.

Using a new animal model, we hypothesised that maternal high fat diet (HF) induced obesity and acute viral infection would each independently induce a pro-inflammatory, pro-oxidative environment in pregnancy, resulting in impaired placental cellular turnover, but these effects would be gestational-age dependent. We further hypothesised that co-exposure to obesity and infection would exacerbate placental stress, leading to greater disruption of normal placental phenotype and fetal growth than the effects of these exposures alone. By understanding how maternal obesity and viral infection interact to alter the placental inflammatory and oxidative stress milieux we can improve understanding of placental response to these common conditions in pregnancy and the mechanisms that drive poor fetal growth, development, and altered health trajectories.

## Methods

### Animal model and study design

All experiments were approved by the animal care committee at Carleton University (AUP 114994). Additionally, we reported procedures and outcomes following the ARRIVE guidelines (Supplementary Table 1)^42^. Five to six weeks old male and female C57BL/6J mice were housed in a single room under standard conditions (25°C and 12:12 light-dark cycle) with free access to food and water. Females were randomized into two main dietary groups **1**. mice fed a HF diet (Teklad custom diet TD.210535 with 62% fat, 4% inulin, 18.8% protein, 19.2% carbohydrates and 5 kcal/g energy density; n=28) or **2**. control diet (Teklad custom diet TD.210534 with 15.8% fat, 4% inulin, 20.9% protein, 63.3% carbohydrates and 3.5 kcal/g energy density; Inotiv, Indianapolis, In, United States; CON, n=34); *ad libitum* six to eight weeks before mating and throughout pregnancy. A subset of dams from each dietary group received a single injection of either 10 mg/kg of Poly(I:C) (a dsRNA viral mimic), or saline as vehicle (0.9%; 0.1 ml) 24 hours before sample collection at gestational day (GD)12.5, 15.5 or 18.5, and creating four experimental groups: CON-VEH, CON-POLY, HF-VEH, HF-POLY per GD (n=5-9/group at each GD).

Inulin, a soluble prebiotic fibre, was added to the diets to address the fact that most purified diets, including high fat diets, are low in fibre, which can alter gut microbiota and metabolic outcomes, confounding interpretation relative to human diets^43,44^. Gestational time points were chosen to investigate exposures throughout key windows of placental maturation: At GD12.5 (early pregnancy) main placental structures are established and can be differentiated^45^; at GD 15.5 (mid-pregnancy) structures are functionally mature^45^; until GD18.5 (late pregnancy), the placenta continues to remodel and grow to optimise placental function^38,45,46^. Poly(I:C) dose was determined based on a pilot study in pregnant C57BL/6J mice, where three doses (2.5, 5.0 and 10 mg/kg) were tested for their ability to elicit a maternal inflammatory response (elevated circulating cytokines) without causing pregnancy loss^47^. Our study confirmed that 10 mg/kg induced maternal immune activation while avoiding pregnancy loss or preterm birth and was selected for subsequent experiments (Supplementary Table 2).

### Biospecimen collection and processing

At each GD, dams were euthanised by cervical dislocation, and trunk blood was collected into heparin-coated tubes following decapitation. Maternal samples, fetuses, and placentae were rapidly dissected from the uterus, weighed, flash frozen in liquid nitrogen and stored at -80°C for molecular analysis, or fixed in 10% neutral-buffered formalin for 48 hours for histological assessments. Tail fragments from all fetuses were collected at the time of harvest to confirm fetal sex by PCR genotyping. DNA was extracted from tail samples using the Extract-N-Amp Tissue PCR Kit (Sigma-Aldrich, Oakville, ON, Canada), according to the manufacturer’s instructions. Genotypes were confirmed by PCR amplification of the target sex chromosomes followed by agarose DNA gel electrophoresis. Primer sequences for genes of interest are listed in Supplementary Table 3. For placental analysis, one male and one female placenta per litter per dietary and infectious group was used for downstream assays.

### Maternal plasma cytokine and chemokine measurements

To determine whether HF diet and Poly(I:C) exposure impacted maternal inflammatory status, pro-inflammatory cytokines or chemokines were quantified in maternal plasma using a commercially available mouse-specific 8-plex assay, (BioRad, Mississauga, ON, Canada). Targets were: interleukin-2 (IL-2), interleukin-4 (IL-4), tumor necrosis factor-alpha (TNF-α), interferon-gamma (IFN-γ), granulocyte-macrophage colony-stimulating factor (GM-CSF), interleukin-5 (IL-5), interleukin-10 (IL-10), IL-1β.

### RNA isolation and expression in the placenta

As previously described^48^, total RNA was extracted from mouse placenta using the RNeasy Mini Kit (CAT #74134; QIAGEN, Toronto, ON, Canada) following manufacturer’s instructions. RNA purity and concentration were assessed by spectrophotometric analysis (Nanodrop) and RNA integrity was verified using gel electrophoresis. Following confirmation of RNA quality, 1 μg RNA was reverse transcribed using 5X iScript Reverse Transcription Supermix (CAT #1725035; BioRad, Mississauga, ON, Canada). A non-reverse transcription (NRT; absence of enzyme) sample was reverse transcribed to provide a negative control for the RT reaction in downstream PCR applications. Real time PCR (Bio-Rad CFX384, Hercules, CA, USA) was used to measure mRNA expression levels of the following inflammatory genes: *Tlr-3, Tlr4, Irf3, Tirap and Il-1β*. Primer sequences for genes of interest and three stably expressed reference genes (*β-actin, Ywhaz, Gapdh*) are listed in Supplementary Table 3. Each qPCR reaction was prepared with SsoAdvanced Universal SYBR Green (CAT #1725270; BioRad, Mississauga, ON, Canada) combined with a 0.8 μM mix of forward and reverse primers specific to each target gene. Standard curves were generated for all genes. All standards, samples, and controls were run in triplicate using the following thermal cycling conditions: initial denaturation at 95°C for 20 seconds, followed by 40 cycles of 95°C for 5 seconds and 60°C for 20 seconds. A melt curve analysis was performed from 65°C to 95°C, increasing by 0.5°C every 5 seconds. qPCR data were analysed using Bio-Rad CFX Manager software (Bio-Rad CFX Maestro 1.1 Version: 4.1.2433.1219). Relative gene expression was quantified by the comparative ΔΔCq method, where the quantification cycle (Cq) of each target gene was normalized to the geometric mean of reference genes (ΔCq). Normalised values were then compared to a designated control group to calculate fold-change expression (ΔΔCq)^49^.

### Immunohistochemistry

To determine whether HF diet and infection affected placental inflammasome expression, lipid peroxidation, and cell proliferation-to-death ratio, 5 μm paraffin embedded placental sections were serially cut from the midline of the placenta from one male and one female placenta per litter, and used for immunohistochemistry to determine target protein localisation and semi-quantitatively assess percent (%) immunoreactive (ir)-area stained for NOD-, LRR- and pyrin domain-containing protein 3 (NLRP3), an inflammasome marker; 4-hydroxynonenal (4-HNE), a marker of lipid peroxidation; and glutathione peroxidase 4 (GPx-4), an antioxidant enzyme; Ki-67, a maker of cell proliferation; and Caspase-3 (non-cleaved, in its inactive form), as a marker of cell death. Primary antibody incubation for NLRP3 (1:100; D4D8T, Cell Signaling Technology; Toronto, ON, Canada), 4-HNE (1:75; ab48506, Abcam; Boston, MA, USA), GPx-4 (1:500; NBP2-76933, Novus Biologicals; Toronto, ON, Canada), Ki-67 (1:200; NCL-L-Ki67-MM1, Leica Biosystems; Concord, ON, Canada) and Caspase-3 (1:1000; ab184787, Abcam; Boston, MA, USA) occurred overnight at 4°C. Next, secondary goat biotinylated anti-mouse antibody (ab64255; Boston, MA, USA) or biotinylated goat anti-rabbit (1:200; Vector BA-1000; Brockville, ON, Canada) in antibody diluent (DAKO; Mississauga, ON, Canada) was applied for 1 hr at RT. For each placenta, three non-overlapping fields of view were randomly captured at 20x magnification (Invitrogen EVOS FL Auto 2 Imaging System; version: 2.0.2094.0, Thermo Fisher Scientific; Ottawa, ON, Canada) within the placental labyrinth zone (LZ: the exchange zone of the mouse placenta^50^) and junctional zone (JZ; the endocrine site of the mouse placenta^50^) under identical exposure settings. Localisation patterns of target protein staining (cell type and distribution in the LZ and JZ) were qualitatively assessed, while semi-quantitative analysis of ir-stained area was performed in ImageJ (version 1.54d) for NLRP3, 4-HNE, GPx-4 and Caspase-3, or positive cell staining for Ki-67. Ir-staining was quantified as the percentage of positively stained area or positive cells relative to the total tissue area within each field of view. Average percent (%) ir-area stained was calculated across the three fields of view for each placenta. Additionally, placental cell proliferation-to-death ratio, an indicator of whether placental cell death exceeds the tissue’s capacity for regeneration, was assessed by quantifying ir proliferating cells (Ki-67-positive cells/nuclei) and apoptotic cells (Caspase-3-positive ir-area stained), expressed as the proportion of positive cells relative to total nuclei within each placental zone. Percentage ir-area stained for each protein was analysed by a single observer blinded to the experimental groups.

### Statistics

All data were tested for normality using the Shapiro-Wilk test. Data that were non-normal were log transformed to achieve normality, where possible. Unequal variances were assessed using Levene’s test. For outcomes analysed with linear mixed models LMMs; described below, model specifications accounted for unequal variance (e.g., litter effects). Difference in maternal weight gain before pregnancy between HF and CON was analysed by t-test. Separately for each GD subgroup (e.g. dams sacrificed at GD 12.5, 15.5 and 18.5), differences between dietary and infectious groups (CON-VEH, CON-POLY, HF-VEH, HF-POLY) for maternal and placental data were analysed using a LMM, considering litter as a random effect, to evaluate the main fixed effects of diet and infection. Placental and fetal data were also analysed stratified by fetal sex. For parametric data, post hoc comparisons between groups were performed using Tukey’s test, following the GLMM analysis. Outcomes with a single independent measurement per dam (e.g., maternal outcomes) were analysed using two-way ANOVA followed by Tukey’s post hoc test. To account for litter effect (i.e. that some fetuses/placentae come from the same litter), outcomes involving multiple fetuses per litter (e.g., placental and fetal data) were analysed using linear mixed models (LMM), with litter included as a random effect. Non-parametric data that could not be log transformed were analysed using Kruskall-Wallis and Steel Dwass post hoc. All data are presented as means ± standard deviation or for non-normal data, median and interquartile range (IQR) with 95% confidence interval diamonds (CI) in figures. Data transformed for analyses are presented as untransformed values. Statistical significance is p≤0.05.

We further reported effect sizes to provide information on the magnitude and potential biological relevance of our findings, in addition to statistically significant effects (p≤0.05), as moderate effect sizes might be biologically relevant. For outcomes analysed with two-way ANOVA, we report effect sizes as partial eta squared (ηp^2^), which quantifies the proportion of variance explained by a given effect relative to error variance. Effect sizes for two-way ANOVA were interpreted according to Cohen’s guidelines: small (0.01), medium (0.06), and large (0.14) effect size^51^. Traditional effect sizes (e.g. Cohen’s d) cannot be used in LLM and with non-parametric data where HL estimates are reported^52^. Accordingly, fixed effect estimates (B coefficients) are reported with standard errors and 95% confidence intervals (CI). Additionally, when maternal or placental data could not be log transformed to achieve normality, we report the Hodges-Lehmann (HL) estimate of the median difference between groups with 95% CI. In these last two cases, effect of biological magnitude was not categorized, as standardized thresholds are not well established. Data were analysed using JMP statistical software (Version:18.0.2).

## Results

### Maternal infection, but not diet, impacts maternal systemic inflammatory profile in early pregnancy

Across gestation, HF diet was not associated with any changes in inflammatory markers in maternal circulation. In contrast, in early pregnancy (GD12.5), but not in mid-(GD15.5) or late pregnancy (GD18.5), maternal infection was associated with an altered systemic inflammatory profile, characterised by increased plasma levels of the anti-inflammatory cytokine IL-10 (ηp^2^=0.30, p=0.02) and a concomitant decrease in the pro-inflammatory cytokine IL-5 compared to vehicle-exposed dams (ηp^2^=0.22, p=0.02) (Figures 1A, D). Additionally in late pregnancy, infection was associated with a significant effect in increasing maternal plasma TNF-α levels compared to VEH (ηp^2^= 0.28, p=0.05; Supplementary Table 4). Co-exposure to HF diet and infection did not further impact maternal systemic levels of pro-inflammatory cytokines at any GD.

**Figure 1.**
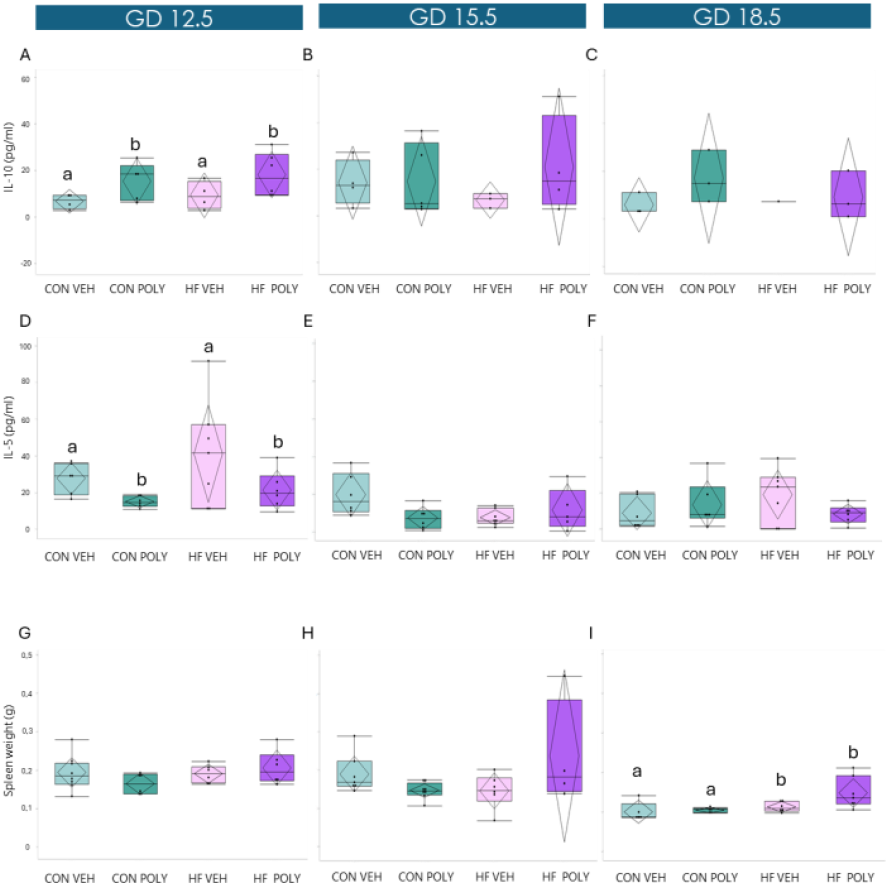
Effect of maternal HF diet and infectious exposure on maternal systemic inflammatory markers and spleen weight across gestation. **A**. Maternal IL-10 plasma levels at GD12.5. Infection was associated with decreased circulating IL-10 levels compared to VEH (p=0.02). **B, C**. Maternal IL-10 plasma levels at GD15.5 and 18.5. **D**. Maternal IL-5 plasma levels at GD12.5. Poly(I:C) exposure was associated with increased circulating IL-10 levels compared to VEH (p=0.02). **E, F**. Maternal IL-5 plasma levels at GD15.5 and 18.5. **G, H**. Maternal spleen weight at GD12.5 and 15.5. **I**. Maternal spleen weight at GD18.5. Maternal HF diet was associated with increased maternal spleen weight compared to CON (p=0.01). Data are quantile box plots with 95% CI diamonds. Two-way ANOVA and Tukey’s post hoc or Kruskall Wallis and Steel-Dwass post hoc. Groups not connected by the same letter are significantly different, p≤0.05. GD = gestational day. CON = control. HF = high fat. VEH = vehicle. POLY = Poly(I:C). CI = confidence interval.

Maternal HF diet was associated with increased maternal spleen weight compared to CON at GD18.5 (HL=0.02, 95% CI [0.006, 0.04], p=0.01; Figure 1I), but not at GD12.5 (Figure 1G) or GD15.5 (Figure 1H). Neither infection alone nor co-exposure to HF diet and infection were associated with changes in spleen weight at any gestational day.

### Maternal diet and infection differentially regulate placental inflammatory signalling pathways across gestation

To confirm placental response to HF diet and Poly(I:C), we measured the expression of placental *Tlr3 and Irf-3, Tlr4* and *Tirap* mRNA expression levels. At GD12.5, there was no effect of HF diet on mRNA expression levels of *Tlr3* and *Irf-3*, which was consistent with expression patterns in mid- and late gestation (Figures 2A, D, G, J). However, at GD12.5, HF diet was associated with decreased *Tlr4* mRNA expression [B=0.6 [−0.001, 0.12]; p=0.05; Figure 2G). By GD18.5, HF diet was associated with increased *Tirap* mRNA expression levels compared to CON-fed pregnancies (B=−0.38 [−0.77, 0.01]; p=0.05; Figure 2L). In contrast, maternal acute infection was associated with increased placental *Tlr3* mRNA expression levels at GD12.5 (B=0.63 [0.18, 1.07]; p=0.009; Figure 2A) and a concomitant decrease in placental *Tlr4* mRNA expression levels (B=−0.65 [−1.30, −0.03]; p=0.04; Figure 2G). Placental *Irf3* and *Tirap* mRNA expression levels were also not affected by acute infection at any gestational age (Figures 2D, E, F, J, K, L). Co-exposure to HF diet and infection had no further impact on the expression levels of any placental inflammatory genes throughout pregnancy.

**Figure 2.**
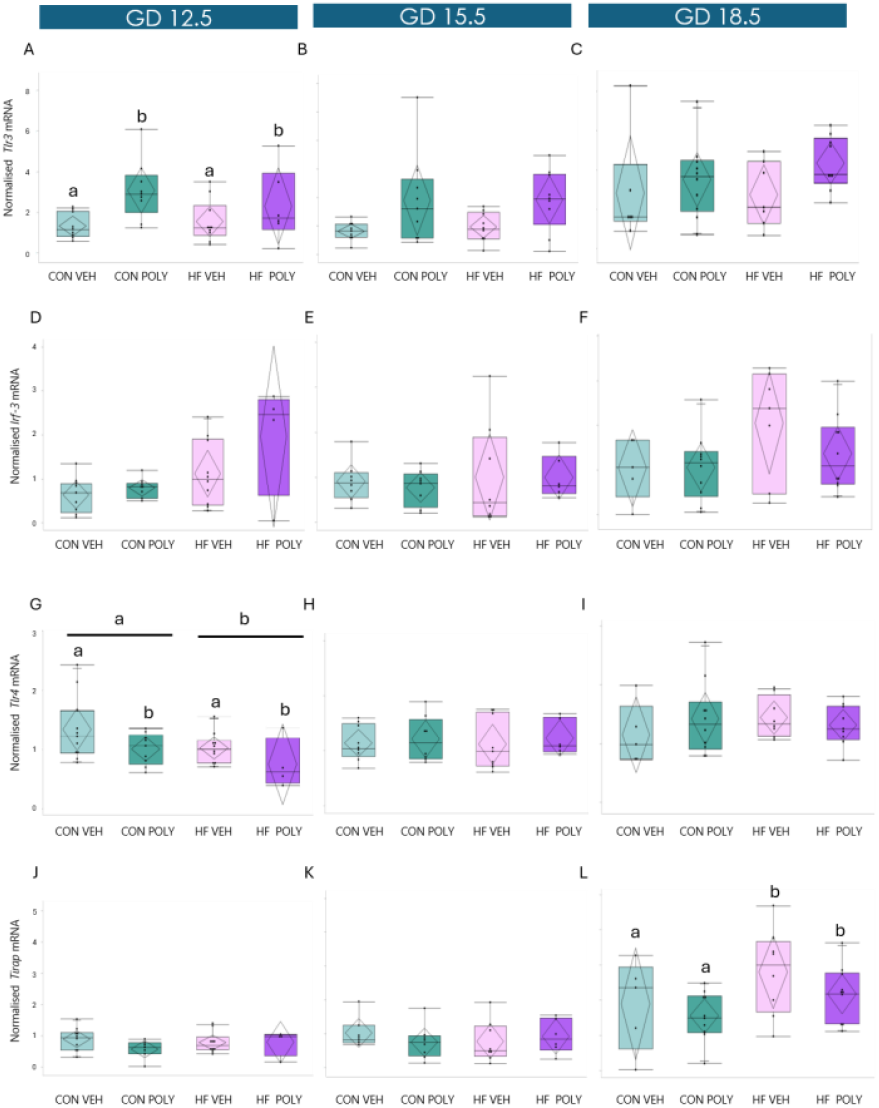
Effect of maternal HF diet and infectious exposure on placental mRNA expression levels of *Tlr*3 and *Tlr4* pathway markers across gestation. **A**. Placental *Tlr3* mRNA expression at GD12.5. Poly(I:C) exposure was associated with increased placental *Tlr3* mRNA expression compared to VEH (p=0.001). **B, C**. Placental *Tlr3* mRNA expression at GD15.5 and 18.5. **D, E, F**. Placental *Irf3* mRNA expression at GD12.5, 15.5 and 18.5. **G**. Maternal HF diet was associated with decreased placental *Tlr4* mRNA expression compared to CON (p=0.05) at GD12.5. Poly(I:C) was associated with decreased placental *Tlr4* mRNA expression compared to VEH (p=0.04) at GD12.5. **H, I**. Placental *Tlr4* mRNA expression at GD15.5 and 18.5. **J, K**. Placental *Tirap* mRNA expression at GD12.5 and 15.5. **L**. Placental *Tirap* mRNA expression at GD18.5. Maternal HF diet was associated with increased placental *Tirap* mRNA expression compared to CON (p=0.05). Data are quantile box plots with 95% CI diamonds. Linear mixed model and Tukey’s post hoc or Kruskal Wallis and Steel-Dwass post hoc. Groups not connected by the same letter are significantly different, p≤0.05. GD = gestational day. CON = control. HF = high fat. VEH = vehicle. POLY = Poly(I:C). CI = confidence interval.

### Maternal diet and infection impact placental lipid peroxidation and antioxidant mechanisms across pregnancy

Given the role of inflammation in oxidative stress, we next investigated whether maternal HF diet and infection influenced placental lipid peroxidation (4-HNE) and antioxidant defence (GPx-4) across gestation. First, ir-staining for 4-HNE and GPX-4 was localised to the JZ and LZ across gestation. ir-4-HNE area stained was mainly expressed in the JZ spongiotrophoblast (SpT) and LZ sysncytiotrophoblast (STB) cytoplasm, while GPx-4 also showed nuclear staining in these placental cell types (Supplementary Figure 2). Second, when observing staining patterns in response to our exposures of interest, we found neither HF diet nor infection alone altered ir-4-HNE or GPx-4 % area stained in the JZ at GD12.5 and 15.5 (Figures 3A, G; 3B, H). However, at GD12.5, co-exposure to HF diet and infection (HF POLY) was associated with decreased ir-4-HNE % area stained in the JZ compared to CON POLY (B=−0.24 [−0.43, −0.06]; p=0.01; Figure 3A). By GD18.5, HF diet was associated with increased placental JZ ir-4-HNE % area stained (B=−0.27 [−0.48, −0.05]; p=0.01; Figure 3C), without changes in ir-GPx-4, whereas infection alone was only associated with increased ir-GPx-4 % area stained (B=−8.46 [−13.51, −3.37]; p=0.004; Figure 3I). Combined HF and infectious exposure There were not associated with no changes in ir-4-HNE and ir-GPx-4 % area stained in the placental JZ at GD18.5 (Figure 3I), in contrast to findings of combined HF diet and infection on lipid peroxidation and antioxidant marker area at GD12.5. Third, in the LZ, HF diet was not associated with changes in placental ir-4-HNE or GPx-4 % area stained at GD12.5 (Figures 3D, J), whereas maternal acute infection was associated with increased lipid peroxidation compared to VEH (B=−0.15 [−0.21, −0.01]; p=0.03; Figure 3D) and reduced ir-GPx-4 % area stained compared to VEH (B=13.47 [0.00, 26.94]; p=0.05; Figure 3J) at this time. Overall, in the LZ at GD12.5, neither HF diet, infection alone, nor co-exposure, impacted ir-4HNE % area stained (Figure 3D), At GD15.5 and GD18.5, neither HF diet, infection, nor co-exposure were associated with changes in ir-4-HNE (Figures 3E, F) or ir-GPx-4 % area stained (Figures 3K, L).

**Figure 3.**
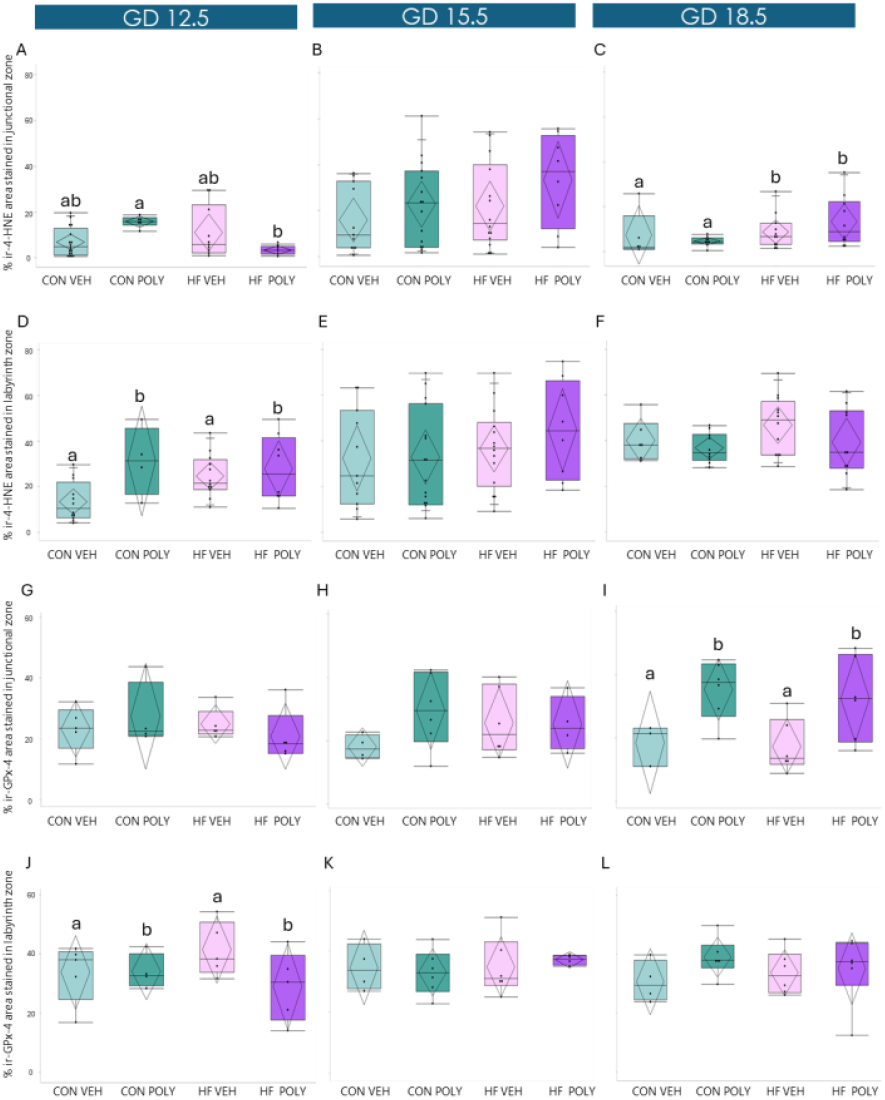
Effect of maternal HF diet and infectious exposure on placental JZ and LZ lipid peroxidation and antioxidant defence across gestation. **A**. Placental JZ lipid peroxidation (% ir-4-HNE area stained) at GD12.5. Co-exposure to HF diet and Poly(I:C) was associated with decreased lipid peroxidation compared to CON POLY (p=0.001). **B**. Placental JZ lipid peroxidation at GD15.5. **C**. Placental JZ lipid peroxidation at GD18.5. Maternal HF diet was associated with increased placental JZ lipid peroxidation compared to CON (p=0.04). **D**. Placental LZ lipid peroxidation at GD12.5. Poly(I:C) was associated with increased placental LZ lipid peroxidation. **E, F**. Placental LZ lipid peroxidation at GD15.5 and 18.5. **G, H**. Placental JZ antioxidant defence (% ir-GPx-4 area stained) at GD12.5 and 15.5. **I**. Placental JZ antioxidant defence at GD18.5. Poly(I:C) exposure was associated with increased JZ antioxidant defence compared to VEH (p=0.04). **J**. Placental LZ antioxidant defence at GD12.5 Placental LZ antioxidant defence. Poly(I:C) exposure was associated with decreased LZ antioxidant defence compared to VEH (p=0.04). **K, L**. Placental LZ antioxidant defence at GD15.5 and 18.5. Data are quantile box plots with 95% CI diamonds. Linear mixed model and Tukey’s post hoc. Groups not connected by the same letter are significantly different, p≤0.05. GD = gestational day. CON = control. HF = high fat. VEH = vehicle. POLY = Poly(I:C). LZ = labyrinth zone. JZ = junctional zone. ir = immunoreactive. CI = confidence interval.

### Maternal diet and infection impact placental inflammation from mid-pregnancy

To determine whether the observed changes in placental oxidative stress were associated with an altered placental inflammatory phenotype, we next measured ir-NLRP3 % area stained in the JZ and LZ. ir-NLRP3 staining in the JZ and LZ SpT and STB occurred in the cell cytoplasm. Further we assessed placental *Il-1β* mRNA expression levels across gestation. At GD12.5, maternal HF diet, infection, or their co-exposure did not significantly affect ir-NLRP3 % area stained in the JZ (Figure 4A). By GD15.5, HF diet was associated with increased ir-NLRP3 % area stained compared to CON (B=−0.13 [−0.25, −0.01]; p=0.03; Figure 4B), whereas infection or co-exposure to HF diet and infection had no further effect. At GD18.5, neither HF diet, infection alone, nor their co-exposure, altered ir-NLRP3 % area stained.

**Figure 4.**
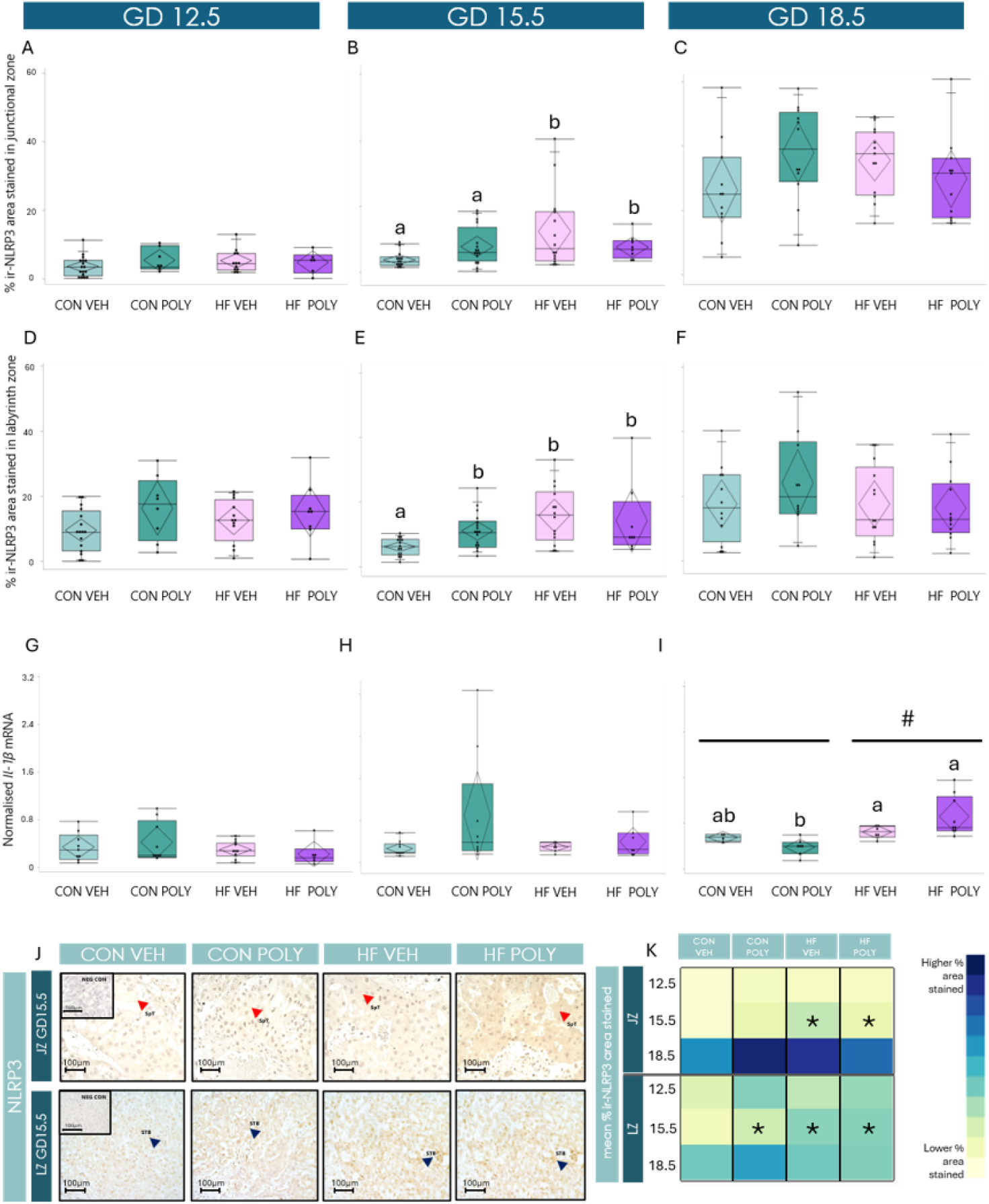
Effect of maternal HF diet and infectious exposure on placental JZ and LZ inflammation across gestation. **A, C**. Placental JZ ir-NLRP3 % area stained at GD12.5 and 18.5. **B**. Placental JZ ir-NLRP3 % area stained at GD15.5. Maternal HF diet was associated with increased placental JZ ir-NLRP3 % area stained compared to CON (p=0.02). **D, F**. Placental LZ ir-NLRP3 % area stained at GD12.5 and 18.5. **E**. Placental LZ ir-NLRP3 % area stained at GD15.5. Co-exposure to HF diet and Poly(I:C) was associated with increased placental LZ ir-NLRP3 % area stained compared to CON VEH (p=0.01). **G, H**. Placental *Il-1β* mRNA expression levels at GD12.5 and 15.5. **I**. Placental *Il-1β* mRNA expression levels at GD18.5. Maternal HF diet was associated with increased placental mRNA expression of *Il-1β* compared to CON (p=0.03). Data are quantile box plots with 95% CI diamonds. Linear mixed model and Tukey’s post hoc. Groups not connected by the same letter are significantly different, p≤0.05 and #indicates significant difference between HF and CON. **J**. Representative images of ir-NLRP3 staining in the placental JZ SpT and LZ STB cytoplasm at GD15.5. Red arrows indicate positive immunostaining in JZ SpT cytoplasm. Blue arrows indicate positive immunostaining in LZ STB cytoplasm. Scale bar = 100μm, magnification = 40x. **K**. Heat map of mean % ir-area stained of NLRP3 by placental zone and GD across exposure groups. Significant differences between dietary and infectious groups in different placental zones and at different GD (p≤0.05) are marked with an asterisk. Darker blue = higher % ir-area stained; lighter blue/yellow = lower % ir-area stained. GD = gestational day. CON = control. HF = high fat. VEH = vehicle. POLY = Poly(I:C). LZ = labyrinth zone. JZ = junctional zone. ir = immunoreactive. SpT = spongiotrophoblast. STB = syncytiotrophoblast. NEG CON = negative control. CI = confidence interval.

At GD12.5, maternal diet alone did not associate with changes in LZ NLRP3 ir-% area stained (Figure 4D). Co-exposure to diet and infection was not associated with changes in ir-NLRP3 % area stained (Figure 4D). At GD15.5, ir-NLRP3 % area stained in the LZ was significantly higher in pregnancies exposed to HF diet and infection compared to CON VEH (B=−0.08 [−0.15, −0.01]; p=0.03; Figure 4E), but not different from the CON POLY or HF VEH groups.

Placental *Il-1β* mRNA expression levels were not altered by HF diet, infection, or their co-exposure in early and mid-pregnancy (Figures 4G, H), however by GD18.5, HF diet was associated with increased placental *Il-1β* mRNA expression levels compared to CON (B=−0.09 [−0.15, −0.03]; p=0.006; Figure 4I). Further, neither infection alone nor co-exposure to HF diet and infection further impacted placental *Il-1β* mRNA expression.

### Maternal HF diet alters placental LZ cell proliferation-to-death ratio in mid- and late gestation

Finally, to determine whether and how HF diet and infection influence placental cellular dynamics, we assessed cell proliferation, cell death and cell proliferation-to-death ratio in the JZ and LZ across gestation. We did not detect ir-Ki-67-positive cells in the JZ SpT nuclei throughout pregnancy, consistent with what has been reported previously^53^. However, ir-Ki-67 staining was observed in the LZ STB nuclei while ir-Caspase-3 staining was observed in the LZ STB cytoplasm. Overall, in the LZ, at GD12.5, neither diet nor infection alone, nor their combined exposure, impacted ir-Ki-67 or ir-Caspase-3 % area stained, or cellular proliferation and death ratio (Figures 5A, D, G). At GD15.5, maternal HF diet was associated with increased placental LZ ir-Ki-67 positive cells compared to CON (B=−0.22 [−0.43, −0.01]; p=0.04; Figure 5B). HF diet was not associated with altered LZ ir-Caspase-3 % area stained (Figure 5E) but was associated with higher LZ cell proliferation-to-death ratio (B=−0.166 [−0.29, −0.03]; p=0.01; Figure 5H). In contrast, infection alone was not associated with changes in ir-Ki-67, Caspase-3 % area-stained levels or cell proliferation-to-death ratio in the LZ at this gestational stage (Figures 5B, E, H). Additionally, neither HF diet nor infection alone, nor their combined exposure, impacted ir-Ki-67 or ir-Caspase-3 % area stained, or cellular proliferation and death ratio (Figures B, E, H). By late pregnancy in the LZ, HF diet was not associated with altered ir-Ki-67 positive cells or cell proliferation-to-death ratio (Figures 5C, I), but was associated with increased ir-Caspase-3 % area stained compared to CON-fed dams (B=−3.44 [−6.23, −0.66]; p=0.01; Figure 5F). Infection or co-exposure to HF diet and infection had no additional effect on placental LZ ir-Ki-67, Caspase-3 % area stained, or cell proliferation-to-death ratio at this stage (Figures 5C, F, I). Placental sex-specific effects between dietary and infectious exposure groups were also observed in the LZ. At GD15.5, exposure to both HF diet and infection was associated with increased ir-Ki-67 positive cells compared to CON POLY in female placentae (p=0.01; Supplementary Figure 4A). By GD18.5, HF diet was associated with increased LZ ir-Caspase-3 % area stained compared to CON in female placentae (p=0.04; Supplementary Figure 4C).

**Figure 5.**
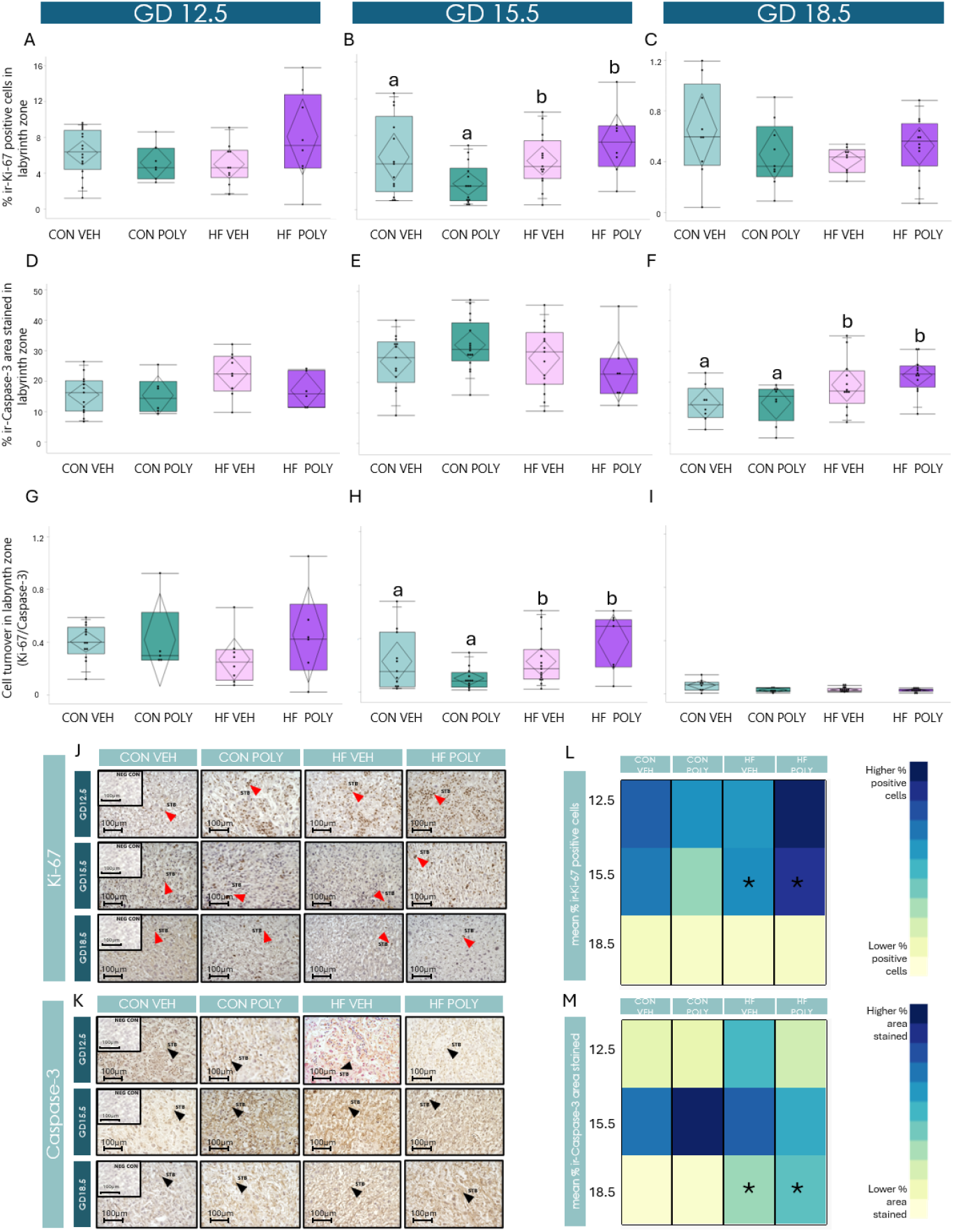
Effect of maternal HF diet and infectious exposure on placental LZ cell proliferation-to-death ratio across gestation. **A, C**. Placental LZ cell proliferation (% ir-Ki-67 positive cells) at GD12.5 and 18.5. **B**. Placental LZ cell proliferation at GD15.5. Maternal HF diet was associated with increased placental LZ cell proliferation compared to CON (p=0.01). **D, E**. Placental LZ cell death (% ir-Caspase-3 area stained) at GD12.5 and 15.5. **F**. Placental LZ cell death at GD18.5. Maternal HF diet was associated with increased placental LZ cell death compared to CON (p=0.002). **G, I**. Placental LZ cell proliferation-to-death ratio (% ir-Ki-67 positive cells/% ir-Caspase3 area stained) at GD12.5 and 18.5. **H**. Placental LZ cell proliferation-to-death ratio at GD15.5. Maternal HF diet was associated with increased placental LZ cell proliferation-to-death ratio compared to CON (p=0.01). Data are quantile box plots with 95% CI diamonds. Linear mixed model and Tukey’s post hoc. Groups not connected by the same letter are significantly, p≤0.05. **J, K**. Representative images of ir-Ki-67 staining in the placental LZ STB nuclei (J) and ir-Caspase-3 staining in the LZ STB cytoplasm (K) across dietary and infectious exposures at different GD. Red arrows indicate positive Ki-67 immunostaining in LZ STB nuclei. Black arrows indicate positive Caspase-3 immunostaining in LZ STB cytoplasm. Scale bar = 100μm, magnification = 40x. **L**. Heat map of mean % ir-Ki-67 positive cells by GD across exposure groups. **M**. Heat map of mean % ir-Caspase-3 area stained by placental zone and gestational day across exposure groups. Significant differences between dietary and infectious groups in different placental zones and at different GD (p≤0.05) are marked with an asterisk. Darker blue = higher % ir-area stained/positive cells; Lighter blue/yellow = lower % ir-area stained/positive cells. CON. GD= gestational day. CON = control. HF = high fat. VEH = vehicle. POLY = Poly(I:C). LZ = labyrinth zone. ir = immunoreactive. SpT = spongiotrophoblast. STB = syncytiotrophoblast. NEG CON = negative control. CI = confidence interval.

## Discussion

Here we investigated the effects of maternal HF diet, infection, and their co-exposure across pregnancy on placental inflammation, oxidative stress, and cellular turnover, given that disruptions in these pathways could explain adverse pregnancy and offspring outcomes seen in populations where the co-exposure to HF diet and viral infection are likely to co-exist. At GD12.5, maternal HF diet alone did not influence placental pro-inflammatory and oxidative pathways. However, infection was associated with immune and oxidative changes, including increased maternal circulating IL-10 and reduced circulating IL-5 levels, increased placental *Tlr3* mRNA expression, greater lipid peroxidation, and reduced antioxidant defence in the LZ. In mid-pregnancy, we showed for the first time that co-exposure of chronic HF diet with acute viral infection was associated with increased LZ ir-NLRP3 % area stained. This pro-inflammatory milieu was accompanied by greater placental LZ cell proliferation and cell proliferation-to-death ratio. By late gestation, maternal HF diet was associated with elevated placental *Tirap* and *Il-1β* mRNA expression levels, JZ lipid peroxidation, and enhanced cell death, reflecting altered cellular proliferation and death ratio.

Overall, while HF diet was gradually associated with increased inflammatory signalling and oxidative stress across gestation, infection alone disrupted maternal and placental inflammatory status and oxidative milieu much earlier in pregnancy, particularly within the LZ, highlighting the distinct timing and zone-specific effects of these exposures. Paradoxically, at GD12.5 maternal infection was associated with increased circulating IL-10 and decreased IL-5 levels, suggesting less inflammation systemically. Recent studies have proposed immunomodulatory effects of type I IFN (a crucial pathway generally activated in response to viral infections) on both innate and adaptive immune cells with capacity to limit inflammation through potent induction of anti-inflammatory molecules, including IL-10^54^. At GD12.5, acute viral infection was also associated with increased placental *Tlr3* mRNA expression levels, consistent with early innate immune activation during a critical period of placental growth and remodelling, probably as a compensatory, gestational age-depended mechanism to prevent exacerbated immune response to viral infective challenges to prevent pregnancy loss^45,55^. Concomitantly, the observed increase in placental *Tlr3* mRNA levels coincided with elevated placental lipid peroxidation and reduced antioxidant defence in the LZ, suggesting that TLR3-mediated signalling may contribute to oxidative stress in early placental development. TLR3 is known to induce ROS production, which is essential for activating inflammatory pathways including nuclear factor kappa-light-chain-enhancer (NF-κB), IRF-3, and signal transducer and activator of transcription 1 (STAT1)^56,57^.

While increases in placental ROS may be locally beneficial to support pathogen clearance^20^, excessive levels trigger lipid peroxidation, producing toxic 4-HNE, which promotes inflammation and cell death^58^. In our model, viral infection was associated with elevated ir-4-HNE % area stained in the LZ, but in the JZ combined HF diet and infection were associated with reduced ir-4-HNE compared to infection alone, suggesting that these placental compartments differ in their vulnerability to oxidative stress. The JZ is a highly metabolic area responsible for endocrine signalling and placental invasion at early developmental stages, constantly producing ROS in response to the metabolic activity of this region^59^. Our results suggest that in the presence of a double burden (HF diet and acute infection) the placenta may upregulate the expression of antioxidant enzymes or adaptive stress responses in a zone-specific manner. The JZ thus is capable of mounting a protective response to increased ROS production through different antioxidant enzymes, which may dampen oxidative insults and preserve placental function^60^. Additionally, infection was associated with decreased LZ ir-GPx4 % area stained, an enzyme responsible for protection against lipid peroxidation^61^, suggesting an increased placental labyrinth vulnerability to lipid peroxidation. Because the LZ is the primary site of maternal-fetal exchange, reduced antioxidant defence could compromise LZ development, ultimately impairing placental efficiency and fetal growth^50^. Overall, these findings suggest that by GD12.5 of gestation, the placenta is more vulnerable to lipid peroxidation than at later stages, and that exposure to infection at this time can weaken placental antioxidant defences, leading to oxidative stress that may impair placental and fetal development as pregnancy progresses.

Our study shows that maternal HF diet modifies the immune response to viral challenge, producing an early anti-inflammatory profile that shifts to a pro-inflammatory state later in gestation. Indeed, at GD15.5, HF diet was associated with increased ir-NLRP3 % area stained in the placental JZ, while in the LZ, exposure to HF diet, infection, or their co-occurrence were each associated with increased ir-NLRP3 % area stained relative to CON VEH, indicating a comparable increase in ir-NLRP3 staining across all exposures. The NLRP3 inflammasome is activated by pathogen-associated RNA and metabolic stress^62^. The observed increase in placental NLRP3 at GD15.5 was not accompanied by increase in *Il-β* mRNA expression. This is consistent with evidence demonstrating that inflammasome activation occurs via a two-step process^63^. First, NLRP3 priming occurs via TLR activation, without necessarily triggering downstream cytokine production^63^. Cytokine production requires subsequent activation (the second activation step) by stress signals, such as potassium ion efflux, lysosomal disruption or mitochondrial dysfunction^63^. In this context, it is possible that chronic exposure to maternal HF diet until mid gestation, and a secondary insult with Poly(I:C), may be enough to start inflammasome priming, while a continued overnutrition ‘stress’ at later gestational stages may be required to fully activate the inflammasome. Therefore, the observed increase in NLRP3 expression at mid gestation may reflect a primed, but not fully activated, inflammatory state in the placenta. Although at this stage *Tlr-3* mRNA % area stained were not upregulated, increased *Tlr3* mRNA and ir-4-HNE levels at earlier GD, might start a signalling cascade that will culminate in increased inflammasome expression at subsequent gestational ages. Indeed, studies have linked increased ROS and lipid peroxidation with NLRP-3 activation^64^. Though less studied in viral infections during pregnancy, NLRP3 activation has been observed in maternal monocytes infiltrating the placenta and producing cytokines like IL-1 and IL-18^65^. Collectively, these findings suggest early dietary and inflammatory exposures may impair placental response to a second burden leading to a worse trajectory for development and function later in gestation.

Overall, maternal HF diet produced a pro-inflammatory placental milieu, which might explain the zone-specific cellular changes we observed. By mid-gestation, HF pregnancies showed higher LZ ir-Ki-67 positive cells despite the normal slowing of proliferation at this stage. This finding is consistent with inflammation-induced regenerative/proliferative signalling^66,67^. These changes in cellular dynamics associated with HF diet exposure persisted until later gestation, but at GD18.5, changes observed were related to increased ir-Caspase-3 area stained. We also observed increased *Tirap* mRNA expression and a further increase of *Il-1β* mRNA expression levels. In parallel, we observed sustained lipid peroxidation in the JZ, possibly induced by the upregulation of *Il-1β* mRNA expression in late pregnancy, which can be associated with elevated ROS^68^. Although late gestation viral infection was associated with a compensatory antioxidant response, this adaptation was likely insufficient to limit JZ lipid peroxidation. Further, while a certain degree of inflammation, oxidative stress, and cell death are physiologically important for initiating labour at term, exaggerated activation of these pathways, particularly in the context of maternal malnutrition, can compromise placental function, potentially impairing fetal health and increasing the risk of poor pregnancy and fetal/offspring outcomes^69^.

Last, we explored whether and to what extent changes in placental development in response to maternal HF diet and acute viral infection were different between male and female placentae. Previous studies in rat models showed that term female placentae have decreased pro-inflammatory cytokines levels and greater adaptive growth compared to male placentae^70,71^. We found that HF diet in mid-pregnancy was associated with decreased *Il-1β* mRNA expression in female, but not male placentae. At the same time, exposure to both HF diet and infection was associated with increased cell proliferation in female placentae. This finding is consistent with the knowledge that female placentae may dampen the inflammatory response under metabolic stress^71^, compared to increased trophoblast production of TNF-α and decreased IL-10 in males^72^, and have higher proliferative capacity under pathological conditions^73^. In contrast by GD18.5, HF diet was associated with increased cell death in female, but not male placentae. Overall, these findings suggest that maternal diet and infection may differentially affect placental inflammation and cell turnover in a sex- and gestational age-dependent manner. Future studies should investigate mechanisms underlying these sex-specific placental responses to maternal HF diet and infection and determine whether these sex-specific placental adaptations drive sex differences in offspring growth and health.

Our study has several strengths. First, our assessment of placental inflammatory, oxidative, and cellular turnover pathways across multiple gestational stages, and in response to two common stressors in pregnancy, reveals for the first time how these systems shift over pregnancy in response to maternal HF diet, acute viral infection, and their combined exposure. Understanding these temporal dynamics is critical for identifying gestational windows when interventions may prevent placental dysfunction and improve pregnancy outcomes. Our model also controlled for fibre-related effects on gut health and systemic inflammation that are not considered in most animal models of maternal obesity/HF diet^41^. Both HF and CON diets contained 4% inulin to maintain adequate fibre levels, ensuring that observed effects reflected dietary fat content rather than confounding effects of fibre deficiency^67−71^. The presence of fibre in our HF diet may explain, at least in part, why our model does not generate an extensive systemic pro-inflammatory response, consistent with the low grade-inflammation produced by obesogenic diets. Further, our Poly(I:C) model reflects an acute rather than chronic viral exposure, which allows us to induce maternal immune activation at defined gestational stages and directly assess its downstream effects on placental development and function and without resulting in fetal/pregnancy loss. Our model also provides insight into how acute viral exposure interacts with the placental inflammation established by maternal obesity. Additionally, we show for the first time that chronic maternal HF diet exposure can compromise the placenta’s capacity to respond to a secondary stressor, emphasising the importance of a comprehensive maternal health assessment in pregnancy, considering coexisting metabolic and infectious diseases, particularly in socially vulnerable populations. Nonetheless, there are some limitations to our work. Our findings are based on cross-sectional analyses at defined gestational stages, which limits our ability to capture dynamic changes in placental inflammatory, oxidative, and cellular responses across gestation. We assessed total (non-cleaved) Caspase-3 ir-expression, which provides information on the abundance of Caspase-3 proteins but does not directly reflect their enzymatic activation status. Future studies should assess its active form^74^, cleaved Caspase-3. Additionally, while inclusion of inulin improves relevance to human dietary patterns, it does not capture the diversity of fibre, fat, and carbohydrate types present in human diets. Finally, as with all mouse models, species differences in placental architecture may limit direct translation to human pregnancy.

Together, our findings show that maternal HF diet and viral infection exert distinct effects on placental pathways across gestation. Earlier in pregnancy, infection primarily activated placental TLR3 signalling and increased lipid peroxidation, reflecting early immune and oxidative activation. In mid-pregnancy, maternal HF diet and infection alone, and in combination, promoted inflammasome activation. In later gestation, HF diet enhanced placental oxidative stress, and disrupted trophoblast cellular dynamics, suggesting maladaptive remodelling of placental tissue. Although maternal HF diet and infection did not exhibit strong interactive effects, our findings suggest that maternal HF diet may prime the placenta for heightened inflammatory and oxidative responses to secondary insults, increasing the risk of impaired placental function and adverse fetal outcomes. By assessing maternal health and placental development at different gestational stages, we demonstrate the importance of considering exposure timing and co-occurrence in shaping maternal-fetoplacental outcomes and identifying critical windows for intervention to reduce the risk of adverse pregnancy and developmental outcomes.

## Data availability

Some or all datasets generated during and/or analysed during the current study are available from the corresponding author on reasonable request.

## Author Contributions

Conceptualisation, methodology: TF, SM, LC, EB, KLC; formal analysis: TF, LC; writing—original draft preparation: TF, KLC; writing—review and editing: TF, LC, EB, KLC; funding acquisition, resources, supervision: KLC.

## Acknowledgments

We would like to acknowledge the funding from the Natural Sciences and Engineering Research Council of Canada (NSERC; RGPIN-2018-05300 and DGECR-2018-00203). Hailey Scott (Department of Health Sciences, Carleton University) for her contribution and assistance throughout the execution and write up of this project, especially in the qPCR methodology and peer review. Erikles Barbosa (Department of Health Sciences, Carleton University) for his contribution during peer review. Israa Zareef, Oluwatomike Aribaloye and Tiffany Kerdilès (Department of Health Sciences, Carleton University) for their assistance during sample collection and pilot study execution.

## Supplementary tables

**Supplementary Table 1.**
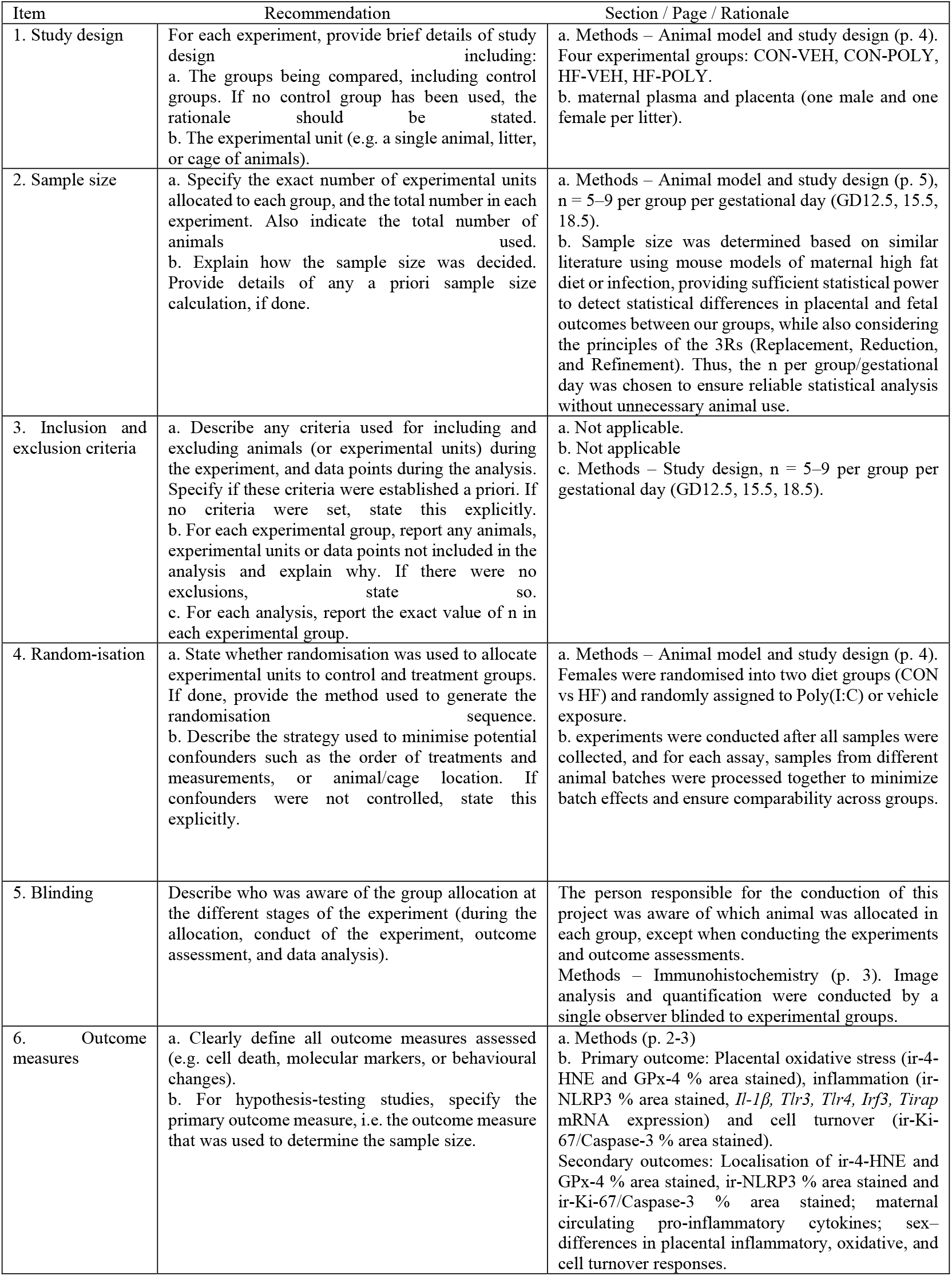
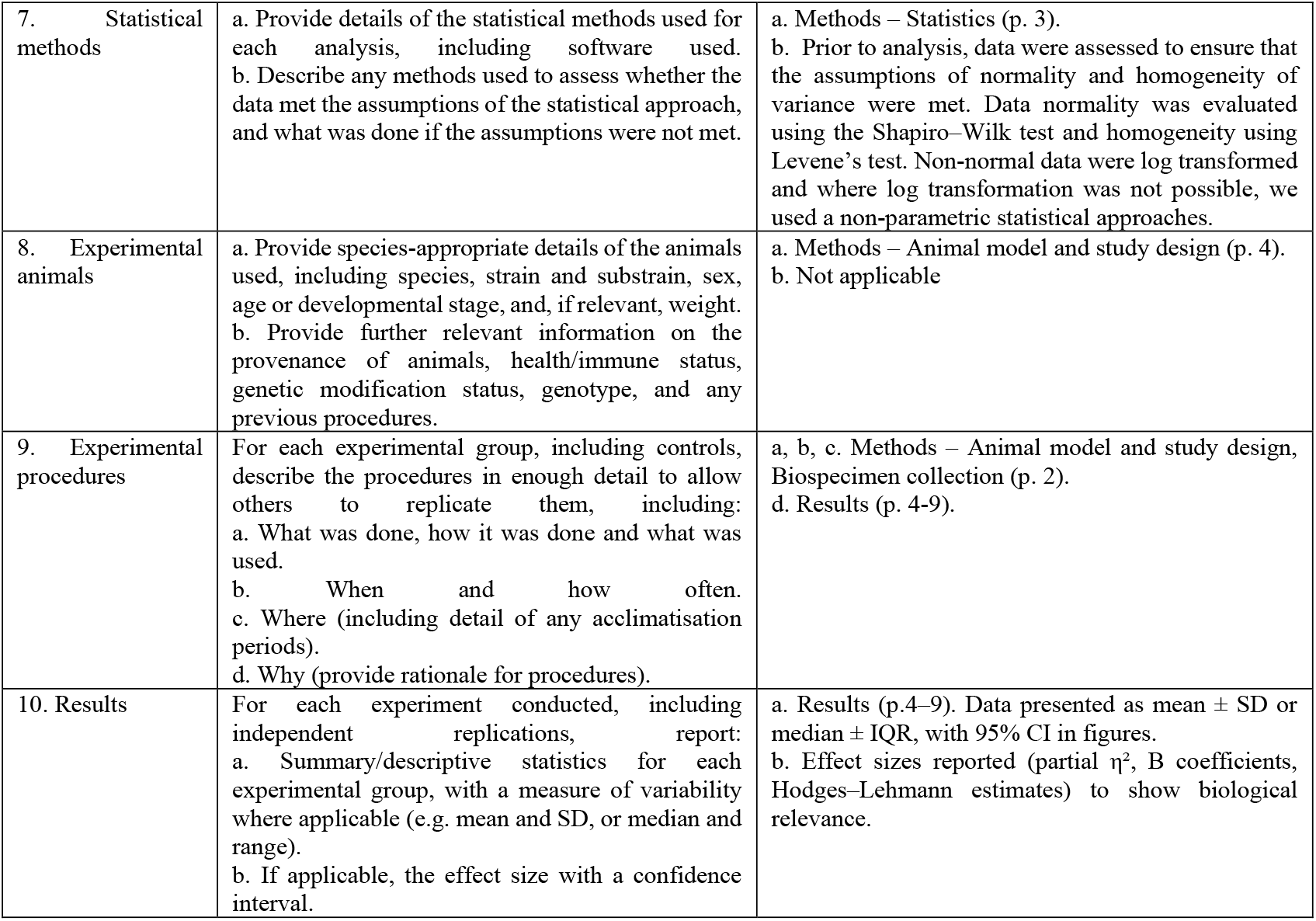
ARRIVE Essential 10 – Compliance Summary.

| Item | Recommendation | Section / Page / Rationale |
| --- | --- | --- |
| 1. Study design | For each experiment, provide brief details of study design including:<br>a. The groups being compared, including control groups. If no control group has been used, the rationale should be stated.<br>b. The experimental unit (e.g. a single animal, litter, or cage of animals). | a. Methods – Animal model and study design (p. 4). Four experimental groups: CON-VEH, CON-POLY, HF-VEH, HF-POLY.<br>b. maternal plasma and placenta (one male and one female per litter). |
| 2. Sample size | a. Specify the exact number of experimental units allocated to each group, and the total number in each experiment. Also indicate the total number of animals used.<br>b. Explain how the sample size was decided. Provide details of any a priori sample size calculation, if done. | a. Methods – Animal model and study design (p. 5), n = 5–9 per group per gestational day (GD12.5, 15.5, 18.5).<br>b. Sample size was determined based on similar literature using mouse models of maternal high fat diet or infection, providing sufficient statistical power to detect statistical differences in placental and fetal outcomes between our groups, while also considering the principles of the 3Rs (Replacement, Reduction, and Refinement). Thus, the n per group/gestational day was chosen to ensure reliable statistical analysis without unnecessary animal use. |
| 3. Inclusion and exclusion criteria | a. Describe any criteria used for including and excluding animals (or experimental units) during the experiment, and data points during the analysis. Specify if these criteria were established a priori. If no criteria were set, state this explicitly.<br>b. For each experimental group, report any animals, experimental units or data points not included in the analysis and explain why. If there were no exclusions, state so.<br>c. For each analysis, report the exact value of n in each experimental group. | a. Not applicable.<br>b. Not applicable<br>c. Methods – Study design, n = 5–9 per group per gestational day (GD12.5, 15.5, 18.5). |
| 4. Random-isation | a. State whether randomisation was used to allocate experimental units to control and treatment groups. If done, provide the method used to generate the randomisation sequence.<br>b. Describe the strategy used to minimise potential confounders such as the order of treatments and measurements, or animal/cage location. If confounders were not controlled, state this explicitly. | a. Methods – Animal model and study design (p. 4). Females were randomised into two diet groups (CON vs HF) and randomly assigned to Poly(I:C) or vehicle exposure.<br>b. experiments were conducted after all samples were collected, and for each assay, samples from different animal batches were processed together to minimize batch effects and ensure comparability across groups. |
| 5. Blinding | Describe who was aware of the group allocation at the different stages of the experiment (during the allocation, conduct of the experiment, outcome assessment, and data analysis). | The person responsible for the conduction of this project was aware of which animal was allocated in each group, except when conducting the experiments and outcome assessments.<br>Methods – Immunohistochemistry (p. 3). Image analysis and quantification were conducted by a single observer blinded to experimental groups. |
| 6. Outcome measures | a. Clearly define all outcome measures assessed (e.g. cell death, molecular markers, or behavioural changes).<br>b. For hypothesis-testing studies, specify the primary outcome measure, i.e. the outcome measure that was used to determine the sample size. | a. Methods (p. 2-3)<br>b. Primary outcome: Placental oxidative stress (ir-4-HNE and GPx-4 % area stained), inflammation (ir-NLRP3 % area stained, *Il-1β*, *Tlr3*, *Tlr4*, *Irf3*, *Tirap* mRNA expression) and cell turnover (ir-Ki-67/Caspase-3 % area stained).<br>Secondary outcomes: Localisation of ir-4-HNE and GPx-4 % area stained, ir-NLRP3 % area stained and ir-Ki-67/Caspase-3 % area stained; maternal circulating pro-inflammatory cytokines; sex-differences in placental inflammatory, oxidative, and cell turnover responses. |
| 7. Statistical methods | a. Provide details of the statistical methods used for each analysis, including software used.<br>b. Describe any methods used to assess whether the data met the assumptions of the statistical approach, and what was done if the assumptions were not met. | a. Methods – Statistics (p. 3).<br>b. Prior to analysis, data were assessed to ensure that the assumptions of normality and homogeneity of variance were met. Data normality was evaluated using the Shapiro–Wilk test and homogeneity using Levene’s test. Non-normal data were log transformed and where log transformation was not possible, we used a non-parametric statistical approaches. |
| 8. Experimental animals | a. Provide species-appropriate details of the animals used, including species, strain and substrain, sex, age or developmental stage, and, if relevant, weight.<br>b. Provide further relevant information on the provenance of animals, health/immune status, genetic modification status, genotype, and any previous procedures. | a. Methods – Animal model and study design (p. 4).<br>b. Not applicable |
| 9. Experimental procedures | For each experimental group, including controls, describe the procedures in enough detail to allow others to replicate them, including:<br>a. What was done, how it was done and what was used.<br>b. When and how often.<br>c. Where (including detail of any acclimatisation periods).<br>d. Why (provide rationale for procedures). | a, b, c. Methods – Animal model and study design, Biospecimen collection (p. 2).<br>d. Results (p. 4-9). |
| 10. Results | For each experiment conducted, including independent replications, report:<br>a. Summary/descriptive statistics for each experimental group, with a measure of variability where applicable (e.g. mean and SD, or median and range).<br>b. If applicable, the effect size with a confidence interval. | a. Results (p.4–9). Data presented as mean ± SD or median ± IQR, with 95% CI in figures.<br>b. Effect sizes reported (partial $\eta^2$, B coefficients, Hodges–Lehmann estimates) to show biological relevance. |

**Supplementary Table 2.** Pilot study maternal and fetal characteristics stratified by different doses of Poly(I:C) in CON and HF pregnancies at GD12.5.

| GD12.5 CON | VEH | 2.5mg/kg | 5mg/kg | 10mg/kg | p-value |
| --- | --- | --- | --- | --- | --- |
| N | (n=3) | (n=3) | (n=3) | (n=3) |  |
| Placental weight (g) | 0.08 ± 0.005<sup>a</sup> | 0.09 ± 0.005<sup>a</sup> | 0.11 ± 0.005<sup>b</sup> | 0.06 ± 0.005<sup>c</sup> | **<0.0001** |
| Fetal weight (g) | 0.10 ± 0.006<sup>a</sup> | 0.125 ± 0.006<sup>a,b</sup> | 0.13 ± 0.006<sup>b</sup> | 0.006 ± 0.006<sup>c</sup> | **<0.0001** |
| Resorption (n) | 1.33 ± 1.11 | 0.66 ± 1.11 | 0.66 ± 1.11 | 2.66 ± 1.11 | 0.55 |
| Litter size (n) | 6.66 ± 0.91 | 7.00 ± 0.91 | 6.66 ± 0.91 | 7.00 ± 0.91 | 0.65 |
| GD12.5 HF | VEH | 2.5mg/kg | 5mg/kg | 10mg/kg | p-value |
| N | (n=3) | (n=3) | (n=3) | (n=3) |  |
| Placental weight (g) | 0.07 ± 0.01 | 0.09 ± 0.02 | 0.08 ± 0.01 | 0.08 ± 0.02 | 0.12 |
| Fetal weight (g) | 0.08 ± 0.05<sup>a,b</sup> | 0.14 ± 0.11<sup>a</sup> | 0.08 ± 0.01<sup>a,b</sup> | 0.08 ± 0.01<sup>b</sup> | **0.01** |
| Resorption (n) | 0.5 ± 1.36 | 1.66 ± 1.11 | 2.00 ± 1.11 | 5.00 ± 1.11 | 0.82 |
| Litter size (n) | 7.00 ± 1.11 | 6.33 ± 0.91 | 6.33 ± 0.91 | 6.66 ± 0.91 | 0.12 |

Data are presented as means ± SEM. One-way ANOVA and Tukey's post hoc or Kruskal-Wallis chi-square test $p \leq 0.05$ . Groups not connected by the same letter are significantly different. GD = gestational day; CON = control; HF = high fat; VEH = vehicle; POLY = Poly(I:C).

**Supplementary Table 3.**
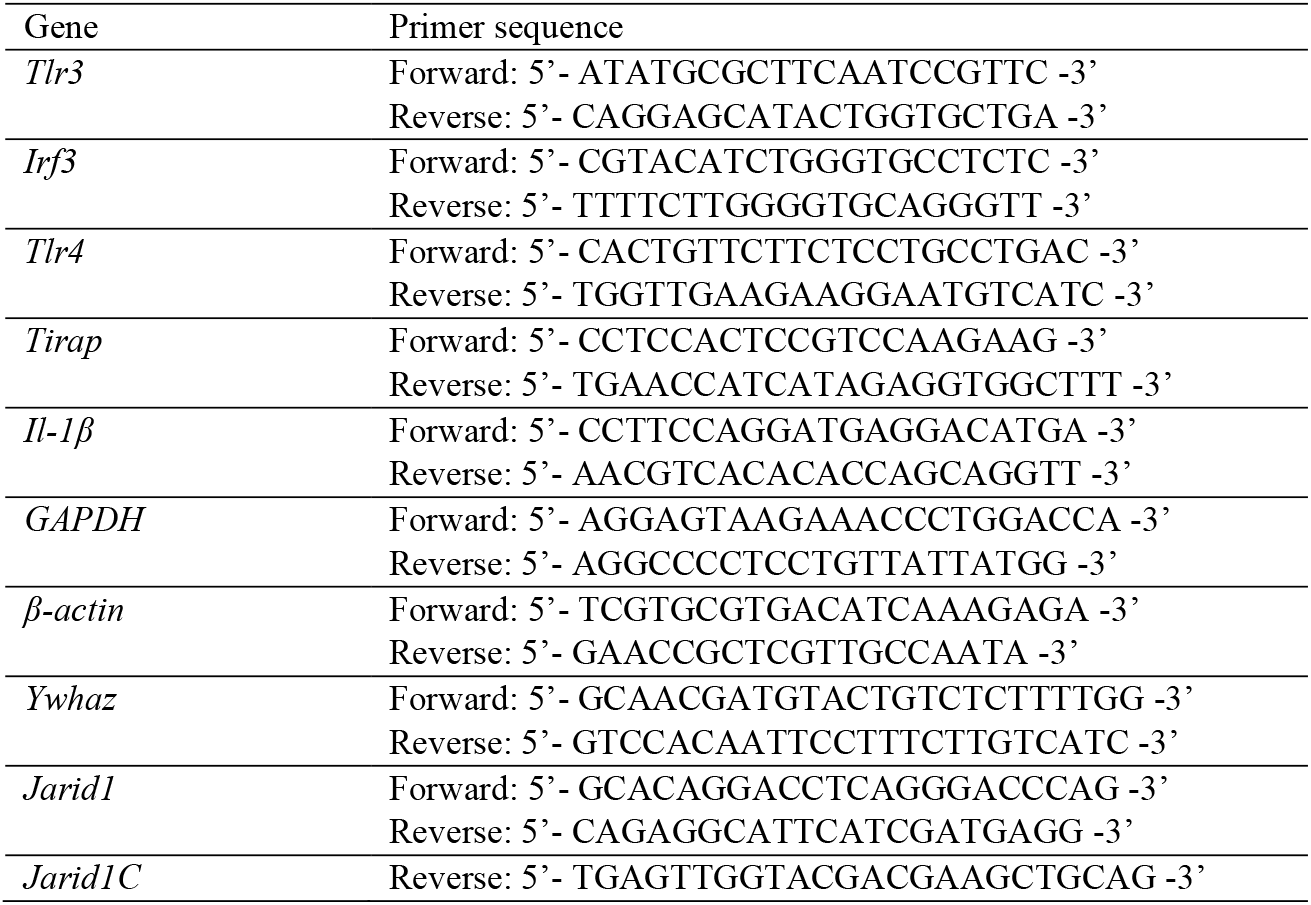
Primer sequences for real-time quantitative PCR gene studies.

**Supplementary Table 4.** Maternal plasma pro-inflammatory cytokines stratified by dietary and infectious exposure at each GD.

| **GD12.5** | **CON VEH** | **CON POLY** | **HF VEH** | **HF POLY** | **p-value** |
| --- | --- | --- | --- | --- | --- |
| N | (n=5) | (n=5) | (n=5) | (n=5) |  |
| IL-10 (pg/ml) | 6.65 ± 3.73<sup>a</sup> | 15 ± 3.33<sup>b</sup> | 9.22 ± 3.73<sup>a</sup> | 18 ± 3.04<sup>b</sup> | **0.02** |
| IL-5 (pg/ml) | 27 ± 6.81<sup>a</sup> | 15 ± 6.81<sup>b</sup> | 41 ± 6.30<sup>a</sup> | 21 ± 6.81<sup>b</sup> | **0.02** |
| TNF-α (pg/ml) | 7.0 [5.46–9.89] | 4.83 [1.90–9.89] | 7.85 [7.41–61.03] | 4.83 [0.76–7.41] | 0.27 |
| IL-1β (pg/ml) | 1.01 ± 0.88 | 0.88 ± 0.72 | 2.36 ± 0.72 | 0.61 ± 0.72 | 0.20 |
| IL-4 (pg/ml) | 0.66 ± 0.15 | 0.59 ± 0.15 | 0.51 ± 0.13 | 0.51 ± 0.15 | 0.80 |
| **GD15.5** | **CON VEH** | **CON POLY** | **HF VEH** | **HF POLY** | **p-value** |
| N | (n=5) | (n=5) | (n=5) | (n=5) |  |
| IL-10 (pg/ml) | 1.11 ± 0.22 | 1.04 ± 0.22 | 0.93 ± 0.20 | 0.78 ± 0.26 | 0.35 |
| IL-5 (pg/ml) | 1.24 ± 0.16 | 0.71 ± 0.16 | 0.82 ± 0.15 | 0.81 ± 0.18 | 0.14 |
| TNF-α (pg/ml) | 6.41 [2.00–8.14] | 2.57 [1.01–4.16] | 2.57 [2.24–3.36] | 2.24 [1.78–2.57] | 0.45 |
| IL-1β (pg/ml) | 0.26 ± 0.52 | 0.63 ± 0.58 | 0.86 ± 0.44 | 0.97 ± 0.58 | 0.81 |
| IL-4 (pg/ml) | 0.18 ± 0.26 | 0.54 ± 0.26 | 0.80 ± 0.26 | 0.62 ± 0.32 | 0.36 |
| **GD18.5** | **CON VEH** | **CON POLY** | **HF VEH** | **HF POLY** | **p-value** |
| N | (n=5) | (n=5) | (n=5) | (n=5) |  |
| IL-10 (pg/ml) | 5.88 ± 5.14 | 17 ± 5.14 | 7.32 ± 8.91 | 9.17 ± 5.14 | 0.49 |
| IL-5 (pg/ml) | 8.8 ± 4.58 | 13 ± 4.58 | 19 ± 4.24 | 8.41 ± 4.58 | 0.10 |
| TNF-α (pg/ml) | 0.42 ± 0.26 | 0.57 ± 0.21 | 0.26 ± 0.23 | 0.13 ± 0.30 | **0.05** |
| IL-1β (pg/ml) | 1.01 ± 0.23 | 0.75 ± 0.18 | 1.09 ± 0.23 | 0.99 ± 0.26 | 0.73 |
| IL-4 (pg/ml) | 0.59 ± 0.17 | 0.55 ± 0.17 | 0.59 ± 0.21 | 0.81 ± 0.21 | 0.53 |

Data are presented as means ± SEM. Data not normally distributed are presented as medians with interquartile range (IQR). Two-way ANOVA and Tukey's post hoc or Kruskal Wallis and Steel Dwass post hoc $p \leq 0.05$ . Groups not connected by the same letter are significantly different. GD = gestational day. CON = control. HF = high fat. VEH = vehicle. POLY = Poly(I:C).

**Supplementary Table 5.** Maternal characteristics stratified by dietary and infectious exposure at each GD.

| Characteristics from dams euthanised at GD12.5 | CON VEH | CON POLY | HF VEH | HF POLY | p-value |
| --- | --- | --- | --- | --- | --- |
| N | (n=9) | (n=6) | (n=7) | (n=6) |  |
| Resorptions (n) | 0.35 ± 0.17 | 1.06 ± 0.20 | 0.82 ± 0.19 | 0.71 ± 0.20 | 0.04 |
| Litter size (n) | 2.12 ± 0.05 | 2.12 ± 0.06 | 2.21 ± 0.06 | 2.13 ± 0.06 | 0.50 |
| Sex ratio (m:f) | 1.40 | 1.71 | 1.08 | 1.14 | 0.87 |
| Characteristics from dams euthanised at GD15.5 | CON VEH | CON POLY | HF VEH | HF POLY | p-value |
| N | (n=9) | (n=10) | (n=8) | (n=5) |  |
| Resorptions (n) | 0.62 ± 0.26 | 0.90 ± 0.23 | 0.28 ± 0.27 | 1.00 ± 0.33 | 0.43 |
| Litter size (n) | 0.90 ± 0.02 | 0.91 ± 0.02 | 0.86 ± 0.03 | 0.94 ± 0.02 | 0.19 |
| Sex ratio (m:f) | 1.14 | 2.00 | 1.40 | 0.67 | 0.42 |
| Characteristics from dams euthanised at GD18.5 | CON VEH | CON POLY | HF VEH | HF POLY | p-value |
| N | (n=9) | (n=7) | (n=8) | (n=8) |  |
| Resorptions (n) | 0.00 ± 0.08<sup>a</sup> | 0.15 ± 0.63<sup>a</sup> | 0.10 ± 0.07<sup>b</sup> | 0.30 ± 0.07<sup>b</sup> | **0.03** |
| Litter size (n) | 0.83 ± 0.08 | 0.93 ± 0.09 | 0.82 ± 0.03 | 0.94 ± 0.02 | 0.09 |
| Sex ratio (m:f) | 1.00 | 0.67 | 2.11 | 1.36 | 0.26 |

Data are presented as means ± SEM. Two-way ANOVA and Tukey's post hoc or Pearson chi-square test $p \leq 0.05$ . Groups not connected by the same letter are significantly different. GD = gestational day; CON = control; HF = high fat; VEH = vehicle; POLY = Poly(I:C) m = Male. f = female.

## Supplementary Figures

**Supplementary Figure 1.**
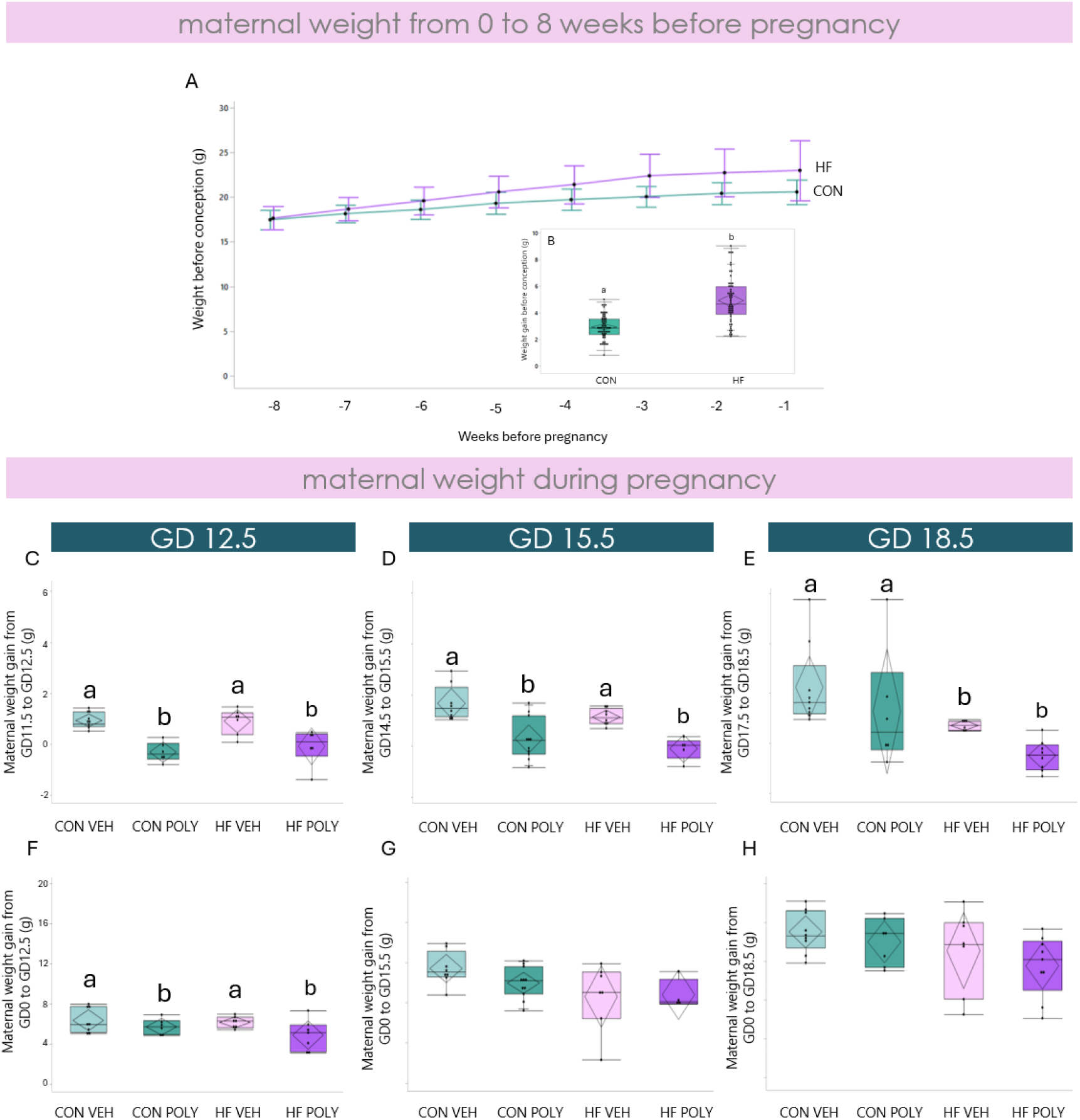
Maternal weight before and during pregnancy in pregnancies exposed to HF diet and infection across ge station. **A**. Maternal weight from 8 weeks before pregnancy until mating in pregnancies exposed to HF or CON diet. **B**. Maternal weight gain from 8 weeks before pregnancy until mating in pregnancies exposed to HF or CON diet. Weight gain was increased in HF-exposed dams compared to CON (p=0.01). **C, D, E**. Maternal weight gain 24h after infectious exposure at GD12.5 (C) GD15.5 (D) and GD18.5. **C**. Poly(:C) was associated with decreased weight gain compared to VEH (p=0.001). **D**. Poly(:C) was associated with decreased weight gain compared to VEH (p=0.001). **E**. Maternal HF diet was associated with decreased weight gain from GD17.5 to 18.5 compared to CON (p=0.02). **F, G, H**. Maternal weight gain from GD0 to GD12.5 (F), GD15.5 (G) and GD18.5 (H). **F**. Poly(:C) was associated with decreased weight gain at GD12.5 compared to VEH (p=0.03). **G, H**. Neither HF diet nor infection impacted maternal weight gain from GD0 to GD15.5 (G) or GD18.5(H). Data are quantile box plots with 95% CI diamonds. t-test and Two-way ANOVA and Turkey’s post hoc. Groups not connected by the same letter are significantly different, p≤0.05. GD = gestational day. CON = control. HF = high fat. VEH = vehicle. POLY = Poly(I:C). CI = confidence interval.

**Supplementary Figure 2.**
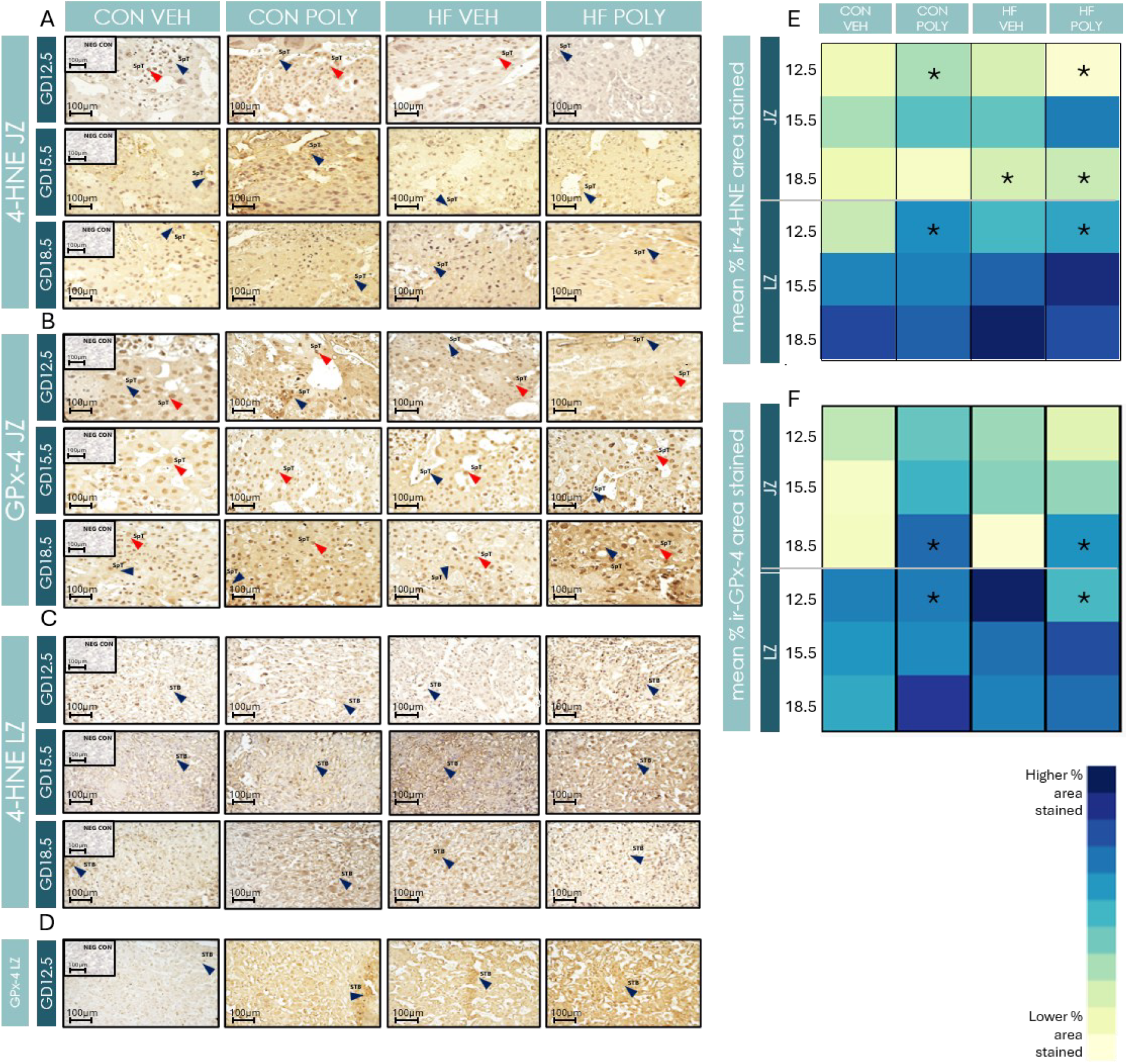
**A**. ir-4-HNE expression (% ir-area stained) in the JZ SpT across dietary and infectious exposures at different GD. **B**. ir-GPx-4 expression (% ir-area stained) in the JZ SpT and STB across dietary and infectious exposures at different GD **C**. ir-4-HNE expression in the LZ STB across dietary and infectious exposures at different GD. **D**. ir-GPx-4 expression in the LZ STB across dietary and infectious exposures at GD12.5. Blue arrows indicate positive immunostaining in JZ SpT and LZ STB. Red arrows indicate positive immunostaining in JZ SpT and LZ STB nuclei. Scale bar = 100μm, magnification = 40x. **E**. Heat map of mean % ir-area stained of 4-HNE by placental zone and GD across exposure groups. **F**. Heat map of mean % ir-area stained of GPx-4 by placental zone and GD across exposure groups. Significant differences between dietary and infectious groups in different placental zones and at different GD (p≤0.05) are marked with an asterisk. Darker blue = higher % ir-expression levels; Lighter blue/beige = lower % ir-expression levels. Intensity of colour indicates higher % ir-expression levels. GD = gestational day; ir = immune reactive; LZ = Labyrinth zone; JZ = Junctional zone; CON = Control; HF = High fat; VEH = Vehicle; POLY = Poly(I:C); 4-HNE = 4-hydroxynonenal; GPx-4 = Glutathione peroxidase-4. SpT = spongiotrophoblast. STB = syncytiotrophoblast.

**Supplementary Figure 3.**
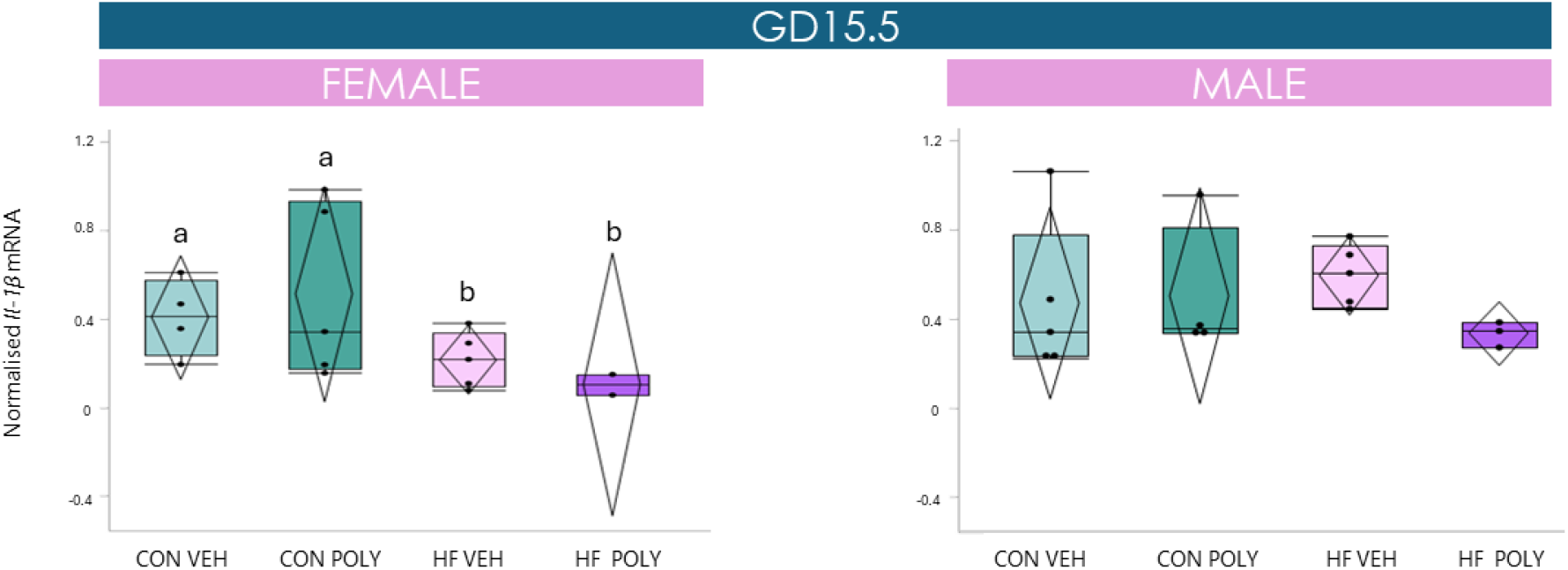
Effect of maternal HF diet and infectious exposure on placental *Il-1β* mRNA expression stratified by sex at GD15.5. **A**. Placental *Il-1β* mRNA expression in female placentae stratified by dietary and infectious group at GD15.5. HF diet was associated with decreased *Il-1β* mRNA expression compared to CON (p=0.02). **B**. Placental *Il-1β* mRNA expression in female placentae stratified by dietary and infectious group at GD15.5. No sex differences were observed in *Il-1β* mRNA expression at GD12.5 or 18.5 and cell death at GD12.5 or 18.5. Data are quantile box plots with 95% CI. Two-way Anova and Tukey’s post hoc. Groups not connected by the same letter are significantly different, p≤0.05. GD = gestational day; CON = control; HF = high fat; VEH = vehicle. POLY = Poly(I:C). CI = confidence interval.

**Supplementary Figure 4.**
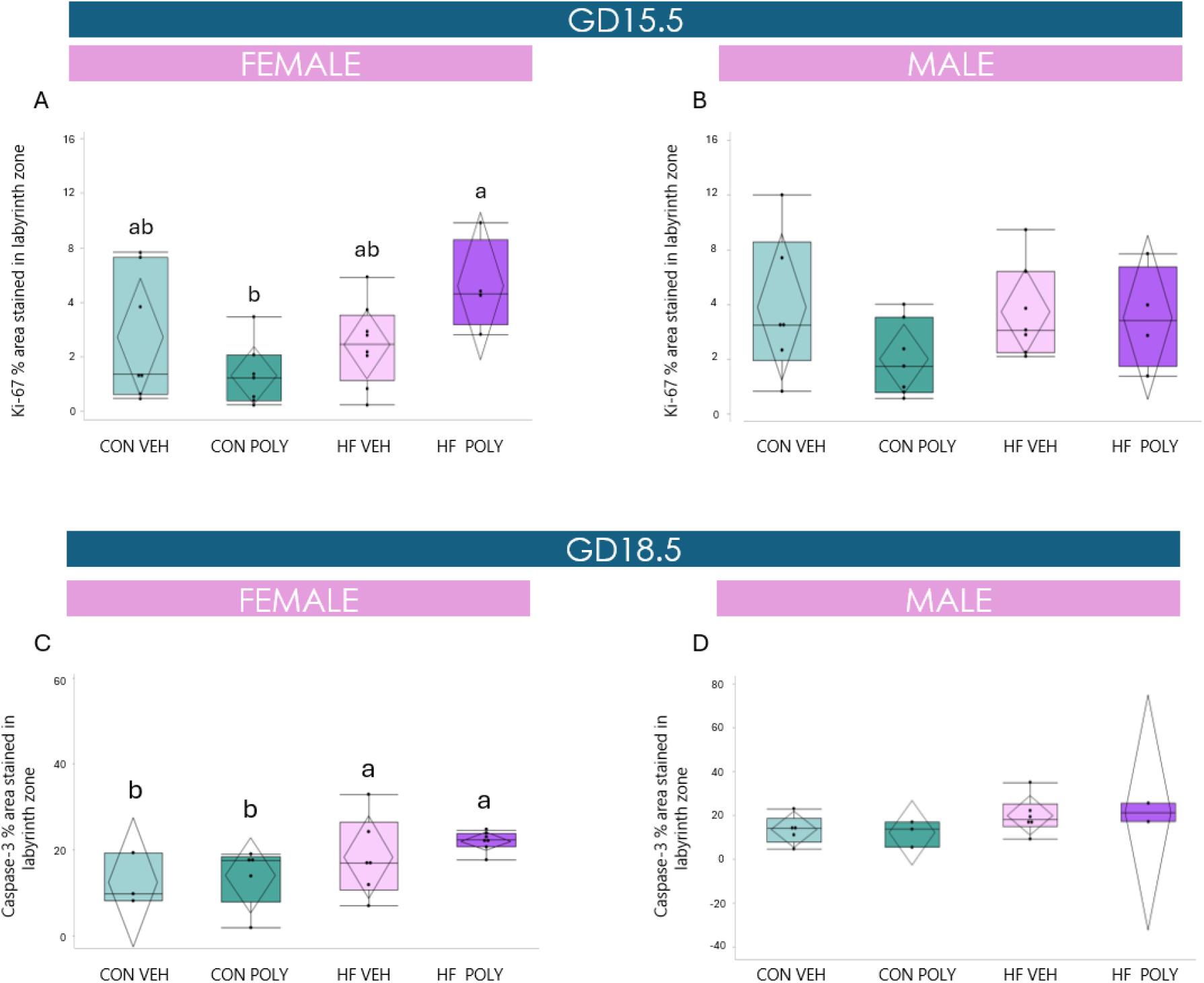
Effect of maternal HF diet and infectious exposure on placental LZ cell proliferation, stratified by sex at GD15.5 and 18.5. **A**. Placental LZ cell proliferation (% ir-Ki-67 area stained) in female placentae at GD15.5. Co-exposure to HF diet and Poly(I:C) was associated with increased placental LZ cell proliferation compared to CON POLY (p=0.01). **B**. Placental LZ cell proliferation in male placentae at GD15.5. **C**. Placental LZ cell death (% ir-Caspase-3 area stained) in female placentae at GD18.5. Maternal HF diet was associated with increased placental LZ cell death compared to CON (p=0.04). **D**. Placental LZ cell death in male placentae at GD18.5. No sex differences were observed in cell proliferation at GD12.5 or 18.5 and cell death at GD12.5 or 18.5. Data are quantile box plots with 95% CI. Two-way Anova and Tukey’s post hoc p≤0.05. Groups not connected by the same letter are significantly different. GD = gestational day; ir = immune reactive; CON = control; HF = high fat; VEH = vehicle. POLY = Poly(I:C). LZ = labyrinth zone; JZ = junctional zone. CI = confidence interval.

## References

1. Ogden CL, Carroll MD, Fryar CD, Flegal KM. Prevalence of Obesity Among Adults and Youth: United States, 2011-2014. NCHS Data Brief. Nov 2015;(219):1–8.

2. Karlsson EA, Marcelin G, Webby RJ, Schultz-Cherry S. Review on the impact of pregnancy and obesity on influenza virus infection. Influenza Other Respir Viruses. Nov 2012;6(6):449–60. doi:10.1111/j.1750-2659.2012.00342.x

3. McClymont E, Albert AY, Alton GD, et al. Association of SARS-CoV-2 Infection During Pregnancy With Maternal and Perinatal Outcomes. JAMA. May 24 2022;327(20):1983–1991. doi:10.1001/jama.2022.5906

4. Amer A, Ayoub A, Brousseau É, Auger N. Risk of severe influenza infection in women with a history of pregnancy complications: A longitudinal cohort study. PLoS One. 2024;19(11):e0313653. doi:10.1371/journal.pone.0313653

5. Group ZW, Medicine CtAoT, (CATMAT) T. Canadian recommendations on the prevention and treatment of Zika virus: Update. Can Commun Dis Rep. May 05 2016;42(5):101–111. doi:10.14745/ccdr.v42i05a01

6. Yakob L. Zika Virus after the Public Health Emergency of International Concern Period, Brazil. Emerg Infect Dis. Apr 2022;28(4):837–840. doi:10.3201/eid2804.211949

7. de Amorin Vilharba BL, Yamamura M, de Azevedo MV, Fernandes WS, Santos-Pinto CDB, de Oliveira EF. Disease burden of congenital Zika virus syndrome in Brazil and its association with socioeconomic data. Sci Rep. Jul 23 2023;13(1):11882. doi:10.1038/s41598-023-38553-4

8. Soma-Pillay P, Nelson-Piercy C, Tolppanen H, Mebazaa A. Physiological changes in pregnancy. Cardiovasc J Afr. 2016;27(2):89–94. doi:10.5830/CVJA-2016-021

9. Racicot K, Mor G. Risks associated with viral infections during pregnancy. J Clin Invest. May 2017;127(5):1591–1599. doi:10.1172/JCI87490

10. Creisher PS, Perry JL, Zhong W, et al. Adverse outcomes in SARS-CoV-2-infected pregnant mice are gestational age-dependent and resolve with antiviral treatment. J Clin Invest. Oct 16 2023;133(20)doi:10.1172/JCI170687

11. Tan L, Lacko LA, Zhou T, et al. Pre-and peri-implantation Zika virus infection impairs fetal development by targeting trophectoderm cells. Nat Commun. Sep 13 2019;10(1):4155. doi:10.1038/s41467-019-12063-2

12. Wang R, Yan W, Du M, Tao L, Liu J. The effect of influenza virus infection on pregnancy outcomes: A systematic review and meta-analysis of cohort studies. Int J Infect Dis. Apr 2021;105:567–578. doi:10.1016/j.ijid.2021.02.095

13. Mulkey SB, Williams ME, Peyton C, et al. Understanding the multidimensional neurodevelopmental outcomes in children after congenital Zika virus exposure. Pediatr Res. Aug 2024;96(3):654–662. doi:10.1038/s41390-024-03056-z

14. Andrade CBV, Lopes LVA, Ortiga-Carvalho TM, Matthews SG, Bloise E. Infection and disruption of placental multidrug resistance (MDR) transporters: Implications for fetal drug exposure. Toxicol Appl Pharmacol. Jan 15 2023;459:116344. doi:10.1016/j.taap.2022.116344

15. Alcalá M, Calderon-Dominguez M, Bustos E, et al. Increased inflammation, oxidative stress and mitochondrial respiration in brown adipose tissue from obese mice. Sci Rep. Nov 22 2017;7(1):16082. doi:10.1038/s41598-017-16463-6

16. Hu Y, He B, Cao Q, et al. Crosstalk of ferroptosis and oxidative stress in infectious diseases. Front Mol Biosci. 2023;10:1315935. doi:10.3389/fmolb.2023.1315935

17. Ferraz T, Benton SJ, Zareef I, Aribaloye O, Bloise E, Connor KL. Impact of Co-Occurrence of Obesity and SARS-CoV-2 Infection during Pregnancy on Placental Pathologies and Adverse Birth Outcomes: A Systematic Review and Narrative Synthesis. Pathogens. Mar 27 2023;12(4)doi:10.3390/pathogens12040524

18. Zemrani B, Gehri M, Masserey E, Knob C, Pellaton R. A hidden side of the COVID-19 pandemic in children: the double burden of undernutrition and overnutrition. Int J Equity Health. 01 2021;20(1):44. doi:10.1186/s12939-021-01390-w

19. de Souza Lima B, Sanches APV, Ferreira MS, de Oliveira JL, Cleal JK, Ignacio-Souza L. Maternal-placental axis and its impact on fetal outcomes, metabolism, and development. Biochim Biophys Acta Mol Basis Dis. Jan 2024;1870(1):166855. doi:10.1016/j.bbadis.2023.166855

20. Denizli M, Capitano ML, Kua KL. Maternal obesity and the impact of associated early-life inflammation on long-term health of offspring. Front Cell Infect Microbiol. 2022;12:940937. doi:10.3389/fcimb.2022.940937

21. Kelly AC, Powell TL, Jansson T. Placental function in maternal obesity. Clin Sci (Lond). 04 2020;134(8):961–984. doi:10.1042/CS20190266

22. Mor G, Cardenas I, Abrahams V, Guller S. Inflammation and pregnancy: the role of the immune system at the implantation site. Ann N Y Acad Sci. Mar 2011;1221:80–7. doi:10.1111/j.1749-6632.2010.05938.x

23. Robson A, Harris LK, Innes BA, et al. Uterine natural killer cells initiate spiral artery remodeling in human pregnancy. FASEB J. Dec 2012;26(12):4876–85. doi:10.1096/fj.12-210310

24. Nardi E, Seidita I, Abati I, et al. The placenta in fetal death: molecular evidence of dysregulation of inflammatory, proliferative, and fetal protective pathways. Am J Obstet Gynecol. Mar 2025;232(3):328.e1-328.e9. doi:10.1016/j.ajog.2024.06.011

25. Mahany EB, Han X, Borges BC, et al. Obesity and High-Fat Diet Induce Distinct Changes in Placental Gene Expression and Pregnancy Outcome. Endocrinology. Apr 01 2018;159(4):1718–1733. doi:10.1210/en.2017-03053

26. Patiño Escarcina JE, da Silva AKCV, Medeiros MOA, et al. Histological Alterations in Placentas of Pregnant Women with SARS-CoV- 2 Infection: A Single-Center Case Series. Pathogens. Sep 26 2023;12(10)doi:10.3390/pathogens12101197

27. Jazwiec PA, Patterson VS, Ribeiro TA, et al. Paternal obesity induces placental hypoxia and sex-specific impairments in placental vascularization and offspring metabolism†. Biol Reprod. Aug 09 2022;107(2):574–589. doi:10.1093/biolre/ioac066

28. Garcez PP, Stolp HB, Sravanam S, et al. Zika virus impairs the development of blood vessels in a mouse model of congenital infection. Sci Rep. Aug 24 2018;8(1):12774. doi:10.1038/s41598-018-31149-3

29. Elgueta D, Murgas P, Riquelme E, Yang G, Cancino GI. Consequences of Viral Infection and Cytokine Production During Pregnancy on Brain Development in Offspring. Front Immunol. 2022;13:816619. doi:10.3389/fimmu.2022.816619

30. Yang X, Li M, Haghiac M, Catalano PM, O’Tierney-Ginn P, Hauguel-de Mouzon S. Causal relationship between obesity-related traits and TLR4-driven responses at the maternal-fetal interface. Diabetologia. Nov 2016;59(11):2459–2466. doi:10.1007/s00125-016-4073-6

31. Wong SW, Kwon MJ, Choi AM, Kim HP, Nakahira K, Hwang DH. Fatty acids modulate Toll-like receptor 4 activation through regulation of receptor dimerization and recruitment into lipid rafts in a reactive oxygen species-dependent manner. J Biol Chem. Oct 02 2009;284(40):27384–92. doi:10.1074/jbc.M109.044065

32. Rajpoot S, Wary KK, Ibbott R, et al. TIRAP in the Mechanism of Inflammation. Front Immunol. 2021;12:697588. doi:10.3389/fimmu.2021.697588

33. England H, Summersgill HR, Edye ME, Rothwell NJ, Brough D. Release of interleukin-1α or interleukin-1β depends on mechanism of cell death. J Biol Chem. Jun 06 2014;289(23):15942–50. doi:10.1074/jbc.M114.557561

34. Li Y, Renner DM, Comar CE, et al. SARS-CoV-2 induces double-stranded RNA-mediated innate immune responses in respiratory epithelial-derived cells and cardiomyocytes. Proc Natl Acad Sci U S A. Apr 20 2021;118(16)doi:10.1073/pnas.2022643118

35. Bao M, Hofsink N, Plösch T. LPS versus Poly I:C model: comparison of long-term effects of bacterial and viral maternal immune activation on the offspring. Am J Physiol Regul Integr Comp Physiol. Feb 01 2022;322(2):R99–R111. doi:10.1152/ajpregu.00087.2021

36. Candia AA, Lean SC, Zhang CXW, et al. Obesogenic Diet in Mice Leads to Inflammation and Oxidative Stress in the Mother in Association with Sex-Specific Changes in Fetal Development, Inflammatory Markers and Placental Transcriptome. Antioxidants (Basel). Mar 28 2024;13(4)doi:10.3390/antiox13040411

37. Ioannidis M, Tjepkema J, Uitbeijerse MRP, van den Bogaart G. Immunomodulatory effects of 4-hydroxynonenal. Redox Biol. Sep 2025;85:103719. doi:10.1016/j.redox.2025.103719

38. Beharier O, Kajiwara K, Sadovsky Y. Ferroptosis, trophoblast lipotoxic damage, and adverse pregnancy outcome. Placenta. May 2021;108:32–38. doi:10.1016/j.placenta.2021.03.007

39. Meihe L, Shan G, Minchao K, et al. The Ferroptosis-NLRP1 Inflammasome: The Vicious Cycle of an Adverse Pregnancy. Front Cell Dev Biol. 2021;9:707959. doi:10.3389/fcell.2021.707959

40. Reddout-Beam C, Hernandez LP, Salak-Johnson JL. Timing of Maternal Stress Differentially Affects Immune and Stress Phenotypes in Progeny. Animals (Basel). Oct 25 2024;14(21)doi:10.3390/ani14213074

41. Louwen F, Kreis NN, Ritter A, Yuan J. Maternal obesity and placental function: impaired maternal-fetal axis. Arch Gynecol Obstet. Jun 2024;309(6):2279–2288. doi:10.1007/s00404-024-07462-w

42. Percie du Sert N, Hurst V, Ahluwalia A, et al. The ARRIVE guidelines 2.0: Updated guidelines for reporting animal research. PLoS Biol. Jul 2020;18(7):e3000410. doi:10.1371/journal.pbio.3000410

43. Pellizzon MA, Ricci MR. Choice of Laboratory Rodent Diet May Confound Data Interpretation and Reproducibility. Curr Dev Nutr. Apr 2020;4(4):nzaa031. doi:10.1093/cdn/nzaa031

44. Zhou P. Effects of prebiotic inulin addition to low- or high-fat diet on maternal metabolic status and neonatal traits of offspring in a pregnant sow model. 2018.

45. Hemberger M, Hanna CW, Dean W. Mechanisms of early placental development in mouse and humans. Nat Rev Genet. Jan 2020;21(1):27–43. doi:10.1038/s41576-019-0169-4

46. Elmore SA, Cochran RZ, Bolon B, et al. Histology Atlas of the Developing Mouse Placenta. Toxicol Pathol. Jan 2022;50(1):60–117. doi:10.1177/01926233211042270

47. Monteiro VRS, Andrade CBV, Gomes HR, et al. Mid-pregnancy poly(I:C) viral mimic disrupts placental ABC transporter expression and leads to long-term offspring motor and cognitive dysfunction. Sci Rep. Jun 17 2022;12(1):10262. doi:10.1038/s41598-022-14248-0

48. Connor KL, Kibschull M, Matysiak-Zablocki E, et al. Maternal malnutrition impacts placental morphology and transporter expression: an origin for poor offspring growth. J Nutr Biochem. Jan 8 2020;78:108329. doi:10.1016/j.jnutbio.2019.108329

49. Livak KJ, Schmittgen TD. Analysis of relative gene expression data using real-time quantitative PCR and the 2(-Delta Delta C(T)) Method. Methods. Dec 2001;25(4):402–8. doi:10.1006/meth.2001.1262

50. Maternal malnutrition impacts placental morphology and transport. An origin for poor offspring growth and vulnerability to disease.

51. Lakens D. Calculating and reporting effect sizes to facilitate cumulative science: a practical primer for t-tests and ANOVAs. Front Psychol. Nov 26 2013;4:863. doi:10.3389/fpsyg.2013.00863

52. Kwak S. Are Only. J Lipid Atheroscler. May 2023;12(2):89–95. doi:10.12997/jla.2023.12.2.89

53. Eaton M, Davies AH, Devine J, et al. Complex patterns of cell growth in the placenta in normal pregnancy and as adaptations to maternal diet restriction. PLoS One. 2020;15(1):e0226735. doi:10.1371/journal.pone.0226735

54. Carlini V, Noonan DM, Abdalalem E, et al. The multifaceted nature of IL-10: regulation, role in immunological homeostasis and its relevance to cancer, COVID-19 and post-COVID conditions. Front Immunol. 2023;14:1161067. doi:10.3389/fimmu.2023.1161067

55. Walker PG, ter Kuile FO, Garske T, Menendez C, Ghani AC. Estimated risk of placental infection and low birthweight attributable to Plasmodium falciparum malaria in Africa in 2010: a modelling study. Lancet Glob Health. Aug 2014;2(8):e460–7. doi:10.1016/S2214-109X(14)70256-6

56. Bassoy EY, Walch M, Martinvalet D. Reactive Oxygen Species: Do They Play a Role in Adaptive Immunity? Front Immunol. 2021;12:755856. doi:10.3389/fimmu.2021.755856

57. Yang CS, Kim JJ, Lee SJ, et al. TLR3-triggered reactive oxygen species contribute to inflammatory responses by activating signal transducer and activator of transcription-1. J Immunol. Jun 15 2013;190(12):6368–77. doi:10.4049/jimmunol.1202574

58. Su LJ, Zhang JH, Gomez H, et al. Reactive Oxygen Species-Induced Lipid Peroxidation in Apoptosis, Autophagy, and Ferroptosis. Oxid Med Cell Longev. 2019;2019:5080843. doi:10.1155/2019/5080843

59. Burdon C, Mann C, Cindrova-Davies T, Ferguson-Smith AC, Burton GJ. Oxidative stress and the induction of cyclooxygenase enzymes and apoptosis in the murine placenta. Placenta. Jul 2007;28(7):724–33. doi:10.1016/j.placenta.2006.12.001

60. Jones ML, Mark PJ, Lewis JL, Mori TA, Keelan JA, Waddell BJ. Antioxidant defenses in the rat placenta in late gestation: increased labyrinthine expression of superoxide dismutases, glutathione peroxidase 3, and uncoupling protein 2. Biol Reprod. Aug 01 2010;83(2):254–60. doi:10.1095/biolreprod.110.083907

61. Martin M, Kumar R, Buchkovich NJ, Norbury CC. HCMV infection downregulates GPX4 and stimulates lipid peroxidation but does not induce ferroptosis. J Virol. Feb 25 2025;99(2):e0185124. doi:10.1128/jvi.01851-24

62. Kelley N, Jeltema D, Duan Y, He Y. The NLRP3 Inflammasome: An Overview of Mechanisms of Activation and Regulation. Int J Mol Sci. Jul 06 2019;20(13)doi:10.3390/ijms20133328

63. Ghonime MG, Shamaa OR, Das S, et al. Inflammasome priming by lipopolysaccharide is dependent upon ERK signaling and proteasome function. J Immunol. Apr 15 2014;192(8):3881–8. doi:10.4049/jimmunol.1301974

64. Zhao S, Chen F, Yin Q, Wang D, Han W, Zhang Y. Reactive Oxygen Species Interact With NLRP3 Inflammasomes and Are Involved in the Inflammation of Sepsis: From Mechanism to Treatment of Progression. Front Physiol. 2020;11:571810. doi:10.3389/fphys.2020.571810

65. Li Y, Qiang R, Cao Z, Wu Q, Wang J, Lyu W. NLRP3 Inflammasomes: Dual Function in Infectious Diseases. J Immunol. Aug 15 2024;213(4):407–417. doi:10.4049/jimmunol.2300745

66. Sferruzzi-Perri AN, Lopez-Tello J, Salazar-Petres E. Placental adaptations supporting fetal growth during normal and adverse gestational environments. Exp Physiol. Mar 2023;108(3):371–397. doi:10.1113/EP090442

67. Yao J, Sterling K, Wang Z, Zhang Y, Song W. The role of inflammasomes in human diseases and their potential as therapeutic targets. Signal Transduct Target Ther. Jan 05 2024;9(1):10. doi:10.1038/s41392-023-01687-y

68. Firmal P, Shah VK, Chattopadhyay S. Insight Into TLR4- Mediated Immunomodulation in Normal Pregnancy and Related Disorders. Front Immunol. 2020;11:807. doi:10.3389/fimmu.2020.00807

69. Sharp AN, Heazell AE, Crocker IP, Mor G. Placental apoptosis in health and disease. Am J Reprod Immunol. Sep 2010;64(3):159–69. doi:10.1111/j.1600-0897.2010.00837.x

70. Phuthong S, Reyes-Hernández CG, Rodríguez-Rodríguez P, et al. Sex Differences in Placental Protein Expression and Efficiency in a Rat Model of Fetal Programming Induced by Maternal Undernutrition. Int J Mol Sci. Dec 28 2020;22(1)doi:10.3390/ijms22010237

71. Braun AE, Mitchel OR, Gonzalez TL, et al. Sex at the interface: the origin and impact of sex differences in the developing human placenta. Biol Sex Differ. Sep 16 2022;13(1):50. doi:10.1186/s13293-022-00459-7

72. Yeganegi M, Watson CS, Martins A, et al. Effect of Lactobacillus rhamnosus GR-1 supernatant and fetal sex on lipopolysaccharide-induced cytokine and prostaglandin-regulating enzymes in human placental trophoblast cells: implications for treatment of bacterial vaginosis and prevention of preterm labor. Am J Obstet Gynecol. May 2009;200(5):532.e1-8. doi:10.1016/j.ajog.2008.12.032

73. Barapatre N, Hansen L, Kampfer C, et al. Trophoblast proliferation is higher in female than in male preeclamptic placentas. Placenta. Dec 2024;158:310–317. doi:10.1016/j.placenta.2024.10.016

74. Wan S, Wang X, Chen W, et al. Exposure to high dose of polystyrene nanoplastics causes trophoblast cell apoptosis and induces miscarriage. Part Fibre Toxicol. Mar 07 2024;21(1):13. doi:10.1186/s12989-024-00574-w

